# The Synaptonemal Complex Central Region Modulates Crossover Pathways and Feedback Control of Meiotic Double-strand Break Formation

**DOI:** 10.1101/2020.07.10.198168

**Authors:** Min-Su Lee, Mika T. Higashide, Hyungseok Choi, Ke Li, Soogil Hong, Kangseok Lee, Akira Shinohara, Miki Shinohara, Keun P. Kim

**Author notes:** These authors contributed equally to this work (Min-Su Lee, Mika T. Higashide).

## Abstract

The synaptonemal complex (SC) is a proteinaceous structure that mediates homolog engagement and genetic recombination during meiosis. Zip-Mer-Msh (ZMM) proteins promote crossover (CO) formation and initiate SC formation. In SC elongation, the SUMOylated SC component Ecm11 and its interacting protein Gmc2 facilitate the polymerization of Zip1, a SC-central region component in budding yeast. Through physical recombination, cytological, and genetic analyses, we here demonstrate that *ecm11* and *gmc2* mutants exhibit chromosome-specific defects in meiotic recombination. CO frequencies were reduced on a short chromosome (chromosome III), whereas CO and non-crossover (NCO) frequencies were increased on a long chromosome (chromosome VII). Further, persistent double-strand breaks (DSBs) occurred in unsynapsed chromosome regions during the late prophase, suggesting the presence of a negative regulation of DSB formation. The Ecm11-Gmc2 (EG) complex could participate in joint molecule (JM) processing and/or double-Holliday junction resolution for CO-designated recombination of the ZMM-dependent pathway. However, absence of the EG complex ameliorated the JM-processing defect in *zmm* mutants, suggesting a role of these proteins in suppression of ZMM-independent recombination. Therefore, the EG complex fosters ZMM-dependent processing and resolution of JMs while suppressing ZMM-independent JM processing and late DSB formation. Hence, EG-mediated SC central regions, which display properties similar to those of liquid crystals, may function as a compartment for sequestering recombination proteins in and out of the process to ensure meiosis specificity during recombination.

## Introduction

During meiosis, pairs of homologous chromosomes (“homologs”) undergo dynamic structural changes and recombination, which is initiated by the formation of programmed DNA double-strand breaks (DSBs). Meiotic recombination is specialized for creating physical connections between homologs, which ensures accurate homologous parental chromosome segregation during the first meiotic division, leading to genetic diversity in a population. Defects in any meiotic recombination process may cause meiotic failure, gamete aneuploidy, and genetic abnormalities (Hunter, 2015).

In many organisms, the formation of meiotic DSBs is catalyzed after meiotic DNA replication at meiotic prophase I by the topoisomerase VI-like protein Spo11 (Lam and Keeney, 2014; Robert et al., 2016). DSB ends subsequently undergo extensive nucleolytic resection to expose a 3’-single-stranded overhang of approximately 800 nucleotides, which is required for homology searching (Cannavo et al., 2013; Garcia et al., 2011; Mimitou et al., 2017). The “first” DSB end recombines and exchanges with a homolog chromatid through a process mediated by the RecA homologs Dmc1 and Rad51, and forms a nascent D-loop that is expanded into the recombination intermediate single-end invasion (SEI) (Cloud et al., 2012; Hong et al., 2013; Lao et al., 2013; Liu et al., 2014; Hong et al., 2019a). The “second” DSB end is thought to engage with the displaced strand of the SEI and produces a double-Holliday junction (dHJ). Interhomolog-dHJs (IH-dHJs) specifically resolve into IH crossover (CO) products; otherwise, the repair of IH non-CO-designated breaks and intermediates via homologs yields non-crossover (NCO) products (Allers and Lichten, 2001; Börner et al., 2004).

Meiotic chromosome axes are organized into a linear array of loops with each pair of tightly conjoined sister chromatids being linked along their entire length to form synaptonemal complexes (SCs) (Page and Hawley, 2004). SCs are meiosis-specific zipper-like proteinaceous structures, comprising axial/lateral elements and a central element that interconnects the axial/lateral elements. A group of proteins known as ZMM proteins (Zip1-3, Spo22/Zip4, Mer3, Msh4, Msh5, and Spo16) initiate SC formation, which is coupled to CO formation (Agarwal and Roeder, 2000; Börner et al., 2004; Chua and Roeder, 1998; Pyatnitskaya et al., 2019; Shinohara et al., 2008; Tsubouchi et al., 2006). ZMM proteins can be classified into three subgroups based on chromosomal and functional criteria (Lynn et al., 2007). Subgroup I includes Mer3 and Msh4-Msh5 (MutSγ), which play a role in diverse DNA repair activities. Subgroup II includes Zip2, Zip3, Zip4/Spo22, and Spo16 (ZZSS), which form the synapsis initiation complex (SIC) to initiate nucleation of the SC. Subgroup III includes Zip1, which contains a coiled-coil domain and a globular domain that correspond to the transverse filament component of the SC. The SC components are involved in reorganization of the recombination complexes (“recombinosomes”) of the SC central region (Lynn et al., 2007). In the absence of ZMM proteins, NCOs occur at high frequencies, whereas CO-designated products and CO formation are strongly defective (Börner et al., 2004). This observation led to the proposal that ZMM proteins are required for the stabilization of recombination intermediates needed to capture the second DSB end into a dHJ, which is then resolved as a CO product. Recombinosomes bind to the regions between the axes and mediate diverse recombination progressions, including homolog partner choice and presynaptic homolog co-alignment in the presence of sister chromatids (Pyatnitskaya et al., 2019). Chromosome axis proteins of the Red1/Mek1/Hop1 complex are required for normal levels of DSBs and for the preferential progression of recombination to form IH recombinants mediated by the RecA homologs Rad51 and Dmc1 (Kim et al., 2010; Hong et al., 2013; Lao et al., 2013). Therefore, axis/SC/recombinosome associations are highly ordered, and homolog pairing persists throughout the CO-fated recombination process (Zickler and Kleckner, 2015).

Once CO/NCO differentiation has occurred in the early prophase, progression to the CO fate involves the production of stable joint molecules (JMs) such as SEI and dHJ intermediates in a ZMM-dependent manner. ZMM-dependent COs, which are often called “type I” COs, exhibit a non-random distribution on chromosomes with positive interference. Some fractions of meiotic DSBs are repaired through a ZMM-independent pathway, which shows random resolution of the Holliday structures yielding both CO and NCO products. The ZMM-independent COs are called “type II” COs and do not show interference. Additionally, dHJs are processed into NCOs through dissolution, which involves the branch migration of HJs. During meiosis, the ZMM-independent pathways seem to be suppressed relative to ZMM-dependent pathways and are thus regulated. However, the molecular nature of this suppression remains unknown.

Multiple feedback controls are capable of downregulating DSBs in order to maintain the proper number and distribution of the DSBs, and thereby managing the recombination events (Keeney et al., 2014; Thacker et al., 2014). The fact that *zmm* mutants demonstrate elevated DSB formation during late meiotic prophase I suggests that homolog engagement regulates the number and distribution of DSB by displacement of Spo11 accessory factors such as Rec114 (Thacker et al., 2014; Anderson et al., 2015; Mu et al., 2020). However, it is not clear whether homolog engagement *per se* or ZMM proteins directly downregulate late meiotic DSB formation.

Small Ubiquitin-like MOdifier (SUMO) plays a role in SC formation (Watts and Hoffmann, 2011). In budding yeast, sumoylation of the SUMO E2-conjugation enzyme Ubc9 is involved in SC assembly and associates with various SC proteins, including the SUMO E3 ligase Zip3 (Cheng et al., 2006; Hooker and Roeder, 2006; Serrentino et al., 2013). Several lines of evidence suggest that SUMOylation of Ecm11, which forms a complex with Gmc2, is important for Zip1 assembly between homologs and that the Ecm11-Gmc2 (EG) complex functions as a component of the SC central region. SUMOylated Ecm11 at early prophase I localizes to the synapsis initiation complex (SIC) in a Gmc2-dependent manner (Humphryes et al., 2013; Voelkel-Meiman et al., 2013). However, the role of SC central regions is less well-defined. Furthermore, SUMOylated Ecm11 localizes to a discrete region of the central element domain that is associated with Zip1 N-termini and limits excess MutSγ-mediated CO formation (Voelkel-Meiman et al., 2015; Voelkel-Meiman et al., 2016). Nevertheless, the underlying functions of EG complex-mediated SC central regions during meiosis remain elusive.

To better define the interplay between homolog engagement and recombination, we further evaluated the regulatory roles of the EG complex in DSB formation and the control of CO-designated DSBs using physical, genetic, and cytological analyses. The results revealed that the EG complex could promote JM processing and/or dHJ resolution for CO-designated recombination. Interestingly, mutation of *ecm11* and *gmc2* resulted in reduced processing of the JMs in the presence of ZMM, whereas *ecm11* and *gmc2* mutants that lacked ZMM were able to effectively process JMs. Moreover, the *ecm11* and *gmc2* deletion mutants showed increased DSB formation, particularly on a long chromosome during late prophase I, suggesting that EG complex-mediated SC polymerization was involved in the feedback control of DSB formation in a chromosome length-dependent manner. Therefore, these results reveal multiple roles for the EG complex in the control of late DSB formation, ZMM-dependent processes that directly regulate type I (interfering) CO-designated DSB repair, and suppression of ZMM-independent recombination (type II, non-interfering COs). We discuss the regulatory role of the EG complex-mediated assembly of the SC central region, which exhibited liquid-crystal properties in these processes.

## Results

### A *gmc2* mutant shows hyper-recombination on chromosome VII

Previous studies have indicated that the EG complex is necessary for efficient Zip1 assembly, which promotes the pairing of a homologous chromosome and SC during meiotic prophase I (Humphryes et al., 2013; Voelkel-Meiman et al., 2013). Furthermore, genetic analysis of *ecm11* and *gmc2* deletion mutants (*ecm11Δ* and *gmc2Δ,* respectively) demonstrated increased CO frequencies within intervals of chromosomes III and VIII of the yeast strain BR1919-8B (Voelkel-Meiman et al., 2016). Therefore, we first analyzed the frequencies of CO and non-CO (non-Mendelian segregation) on chromosomes III and VII in an SK1 background (Higashide and Shinohara, 2016), which revealed synchronous meiosis, as well as on chromosome V in a congenic background (Figure 1A). Consistent with the findings of Voelkel-Meiman et al. (2016), *gmc2Δ* exhibited elevated CO frequencies in four intervals of chromosome VII and in one interval of chromosome III (Figure 1B). In contrast, CO frequencies on two intervals of chromosome III and on one interval of chromosome V in the *gmc2Δ* mutant were similar to those in the wild-type (WT) yeast strain (Figure 1B). One interval of chromosome III and two intervals of chromosome V showed a slight reduction in CO frequencies relative to that in the WT. Most of the loci of the three chromosomes showed more or less increased frequencies of non-Mendelian segregation (Figure 1D). The increase in CO frequencies on the loci of chromosome VII in the *gmc2Δ* mutant was more prominent than that of chromosomes III and V (Figure 1C). When CO interference was examined using nonparental ditype (NPD) ratios, all intervals in the *gmc2Δ* mutant showed decreased CO interference (increased NPD ratios) relative to those in the WT control (Supplemental Figure 1). These results confirmed that the EG complex plays a role in regulating the frequencies and distribution of COs, which may be specific to chromosome properties such as a length.

**Figure 1.**
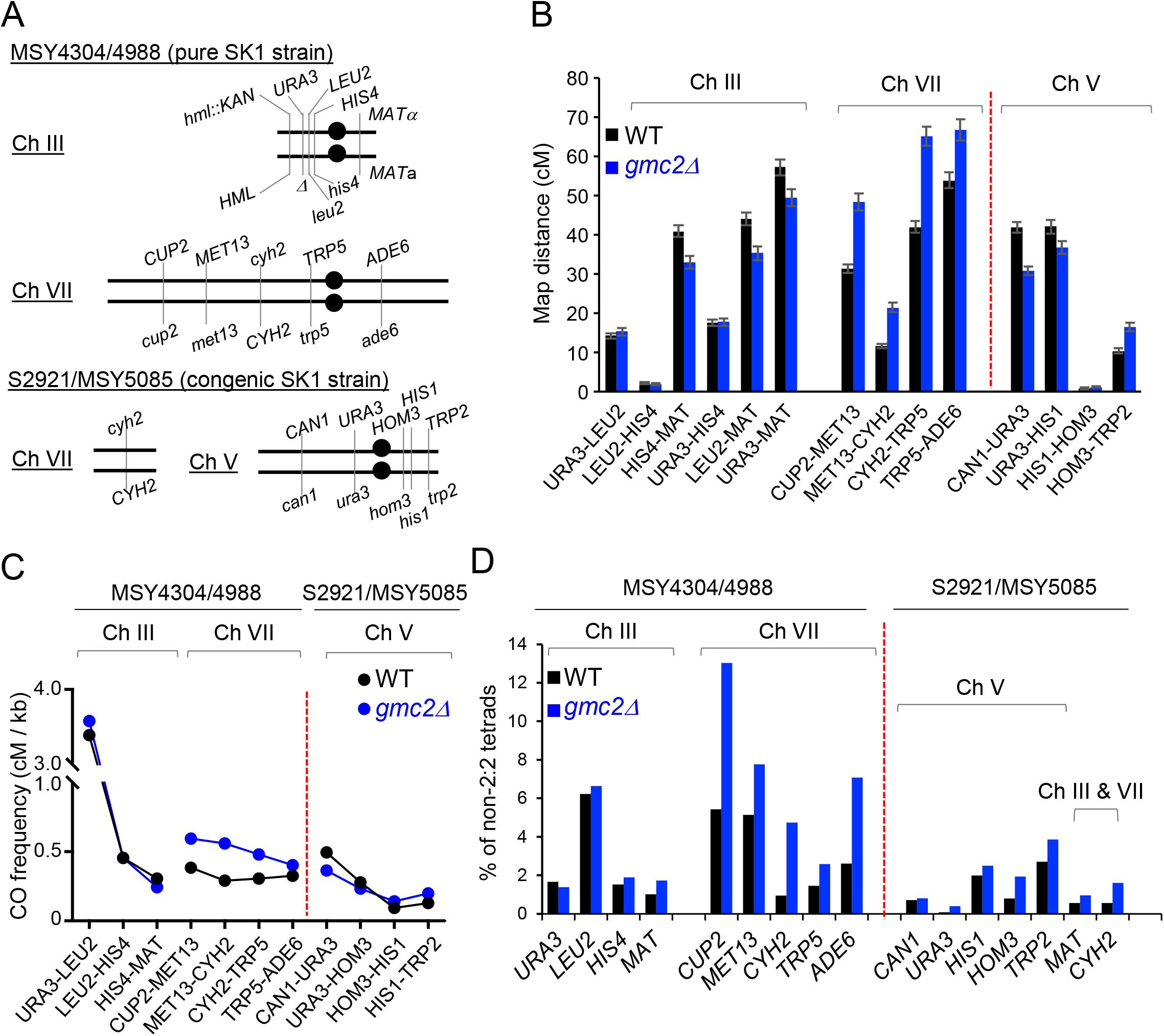
Ecm11–Gmc2 complex regulates meiotic recombination in a bivalent-dependent manner. (A) Schematic representation of the location of marker genes of chromosomes III and VII in the MSY4304/4245 diploid and of chromosomes VII and V in the S2921/MSY5085 diploid. (B) Map distances within each indicated genetic interval of chromosomes III, VII, and V in WT (black) and *gmc2Δ* (blue) strains analyzed using Perkins formula. Error bars show the standard error (S.E.). (C) CO frequencies (cM) per physical length (kb) of each genetic interval of chromosomes III, VII, and V in WT and *gmc2Δ* strains. (D) Frequencies of non-Mendelian segregation of indicated genetic loci in tetrads of WT and *gmc2Δ* strains.

### The EG complex is not required for DSB formation but is necessary for CO-specific recombination at *HIS4LEU2*

#### CO-specific defect

To further investigate the role of the EG complex in meiotic recombination, we used *ecm11Δ/gmc2Δ* single and double mutant strains to analyze recombination intermediates and outcomes at the *HIS4LEU2* locus on chromosome III, which contains a well-controlled single DSB site (Figure 2A). The *ecm11Δ* and *gmc2Δ* mutants showed substantial delays in meiotic division progression by approximately 2 h with approximately 85% of the cells undergoing meiosis (Supplemental Figure 2A). Moreover, the resultant tetrads yielded high levels of four-viable tetrads, 84% for *ecm11Δ,* 88% for *gmc2Δ,* and 84% for *ecm11Δ gmc2Δ* (Supplemental Figure 2B). This finding was consistent with previous results (Humphryes et al., 2013; Voelkel-Meiman et al., 2016). Cell samples of synchronized meiosis cultures were collected at selected time points and subjected to physical analysis for recombination. *XhoI* restriction-site polymorphisms in the *HIS4LEU2* hotspot on chromosome III produced DNA species for DSBs, SEIs, dHJs, and CO products (Figure 2 and Figure 3) as previously described (Oh et al., 2009; Kim et al., 2010; Börner et al., 2004; Hong et al., 2019b). DSBs and COs were evaluated using one-dimensional gel electrophoresis followed by Southern hybridization. COs and NCOs were distinguished by gene conversion of *BamHI* and *NgoMIV* restriction enzyme sites inserted close to the DSB sites at the *HIS4LEU2* locus (Figure 2B, 2D, and 2E). For all physical analyses, radiolabeled probes were used to detect hybridized DNA species.

**Figure 2.**
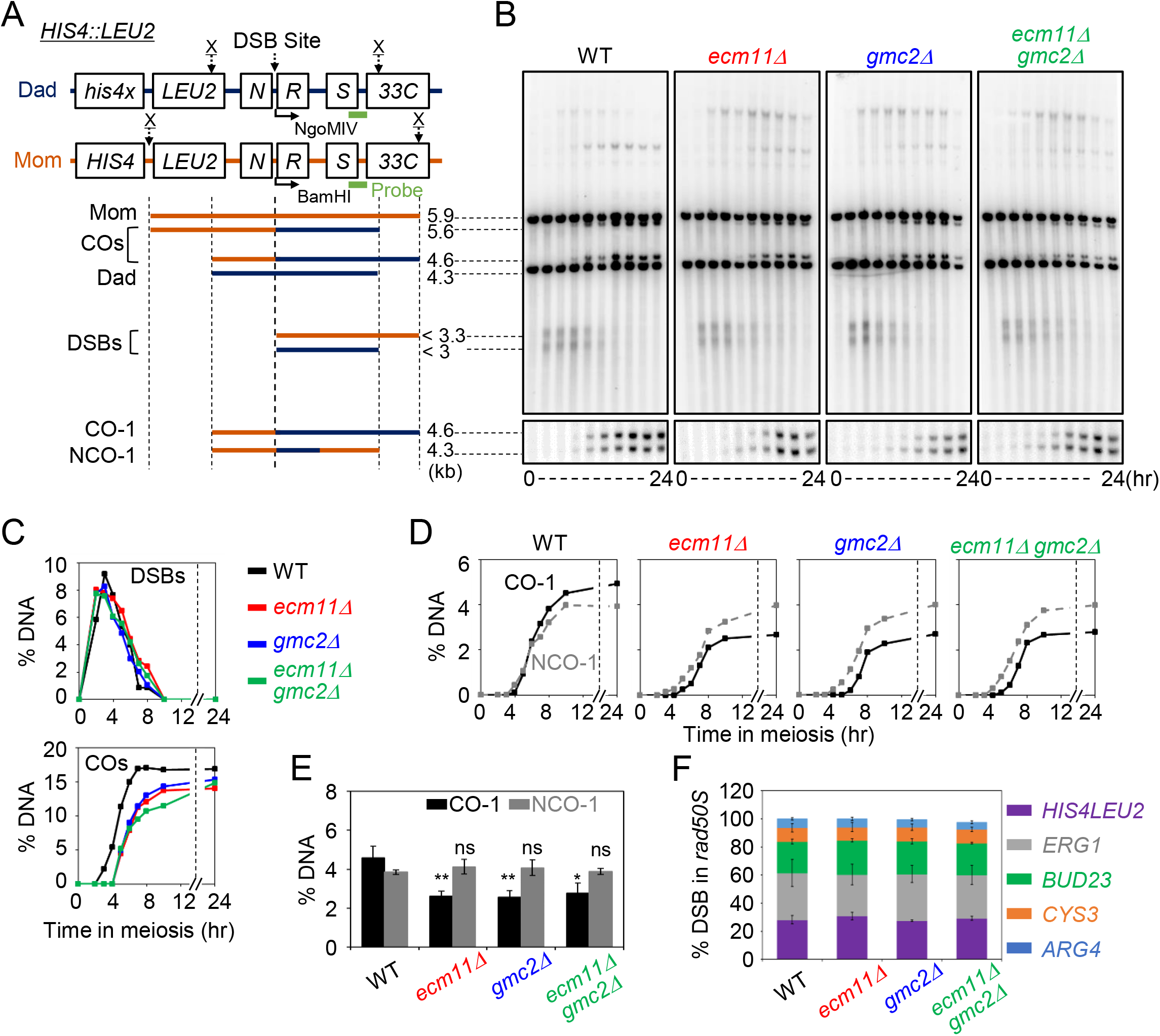
Physical analysis of meiotic recombination in *ecm11Δ* and *gmc2Δ* mutants. (A) Physical map of the *HIS4LEU2* locus of chromosome III showing the *XhoI* (X) restriction endonuclease site and position of the probes for Southern hybridization. Maternal and paternal fragments were distinguished by *Xho*I polymorphisms. For analysis of CO and NCO, DNA was digested with both *Xho*I and *Ngo*MIV endonucleases. Mom, maternal species (5.9 kb); Dad, paternal species (4.3 kb); COs, crossovers (5.6 and 4.6 kb); DSBs, double-strand breaks (<3.3 and <3 kb); CO, crossover (4.6 kb); NCO, non-crossover (4.3 kb). (B) One-dimensional (1D) gel analysis of DSBs, COs, and NCOs in WT, *ecm11Δ, gmc2Δ,* and *ecm11Δ gmc2Δ* strains. Gel analysis (1D) showing Mom, Dad, DSBs, and CO species (top). CO and NCO of recombinants are displayed in the CO/NCO gel analysis (bottom). (C) Quantitation of DSBs and COs shown in panel B. (D) Quantitative analysis of CO (black line) and NCO (gray dashed line). (E) Quantitative analysis of CO and NCO from three independent meiotic time-course experiments (mean ± SD; *N* = 3). Significant differences were analyzed using unpaired *t-* tests (** *p* < 0.01; ns, not significant). (F) Quantitative analysis of DSBs at various loci in *rad50S, rad50S ecm11Δ, rad50S gmc2Δ,* and *rad50S ecm11Δ gmc2Δ* strains. Data indicate mean ± SD (*N* = 3). See Supplementary Figure 4 for more detail.

**Figure 3.**
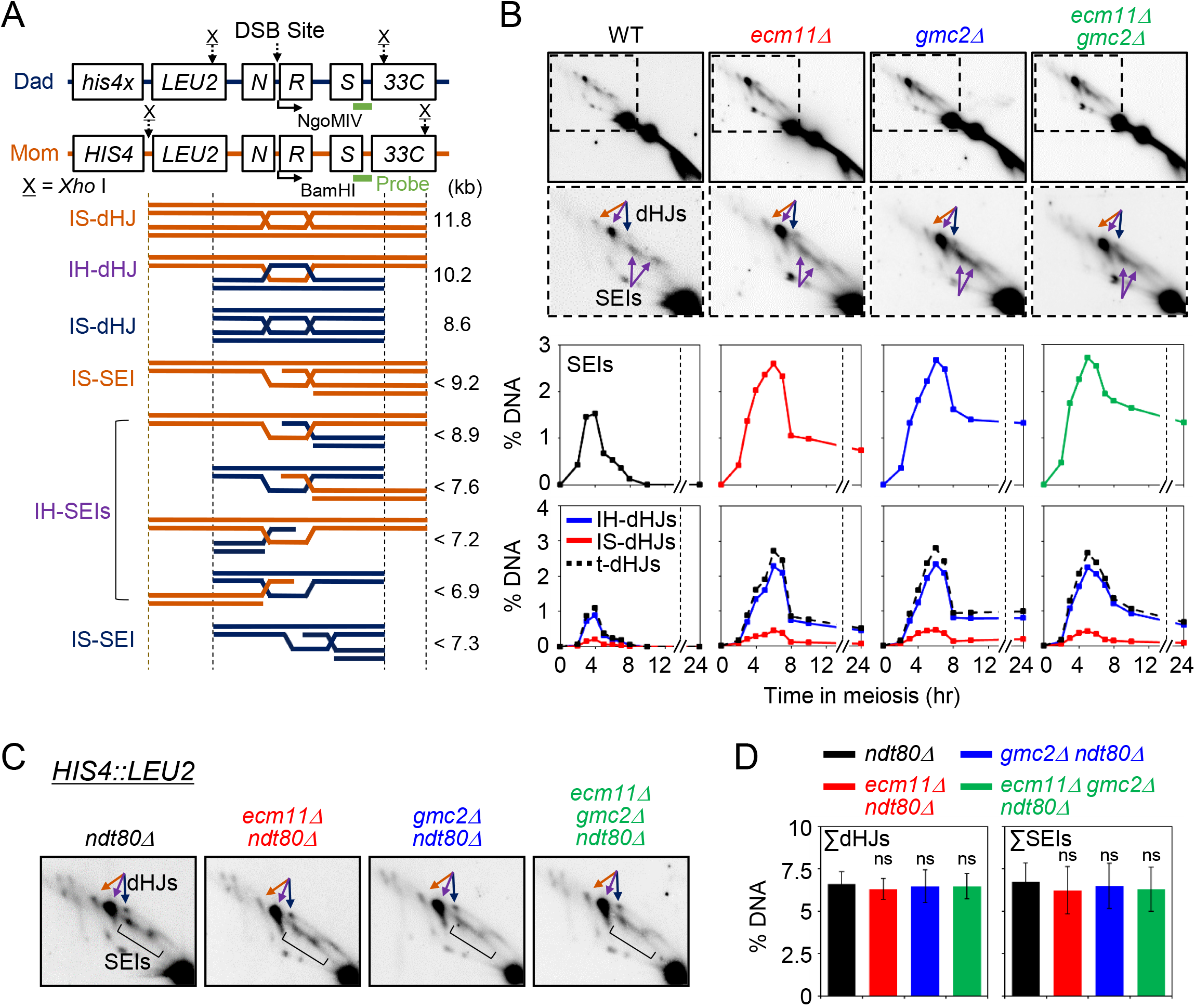
Two-dimensional (2D) gel analysis for *ecm11Δ, gmc2Δ,* and *ecm11Δ gmc2Δ* in WT and *ndt80Δ* backgrounds. (A) Physical map of the *HIS4LEU2* locus. IH-dHJ, interhomolog double-Holliday junction; IS-dHJ, intersister double-Holliday junction; SEIs, single-end invasions. (B) Representative images of 2D analysis of WT, *ecm11Δ, gmc2Δ,* and *ecm11Δ gmc2Δ* strains (top). Quantitation of SEIs and dHJs (bottom). (C) Representative 2D analysis images of the *HIS4LEU2* locus in *ndt80Δ, ndt80Δ ecm11Δ, ndt80Δ gmc2Δ, and ndt80Δ ecm11Δ gmc2Δ* strains. (D) Quantitative analysis of dHJs and SEIs in an *ndt80Δ* background at the *HIS4LEU2* locus. Data indicate mean ± SD (*N* = 3). Significant differences were analyzed using unpaired *t*-tests (ns, not significant).

In the WT species, DSBs appeared and disappeared, followed by the formation of CO products. DSBs in the WT peaked at 3 h and were eventually processed by 8 h (Figure 2B and 2C). The frequency of occurrence of COs (CO-I) and NCOs (NCO-I) was approximately 5% and 4%, respectively (Figure 2D and 2E). The kinetics of DSBs were very similar between the WT and *ecm11Δ /gmc2Δ* strains with respect to the timing as well as maximum levels (Figure 2C). The *ecm11Δ* and *gmc2Δ* single and *ecm11Δ gmc2Δ* double mutants all formed COs (COs in Figure 2C; Figure 4C; Supplementary Figure 3) at 14.9 ± 1.9%, 13.6 ± 1.4%, and 13.7 ± 1.9%, respectively, whereas COs occurred at a frequency of 16.8 ± 1.8% in the WT strain. This indicated a modest reduction in CO frequencies in the *ecm11Δ* and *gmc2Δ* mutant strains. However, total NCO levels were similar to those of the WT strain (~8.7%) (Supplementary Figure 3). Moreover, CO formation in the mutants exhibited an approximate 2-h delay relative to that of the WT strain. In contrast, NCO formation in the mutants occurred with similar timing to that of the WT, suggesting that CO formation was uncoupled from NCO formation in the mutants (Figure 2D). Therefore, in the *ecm11Δ* and *gmc2Δ* mutants, meiotic DSBs at *HIS4LEU2* formed at WT levels with normal post-DSB progression, CO-fated DSB repair suffered from aberrant defects, and NCO formation progressed normally.

**Figure 4.**
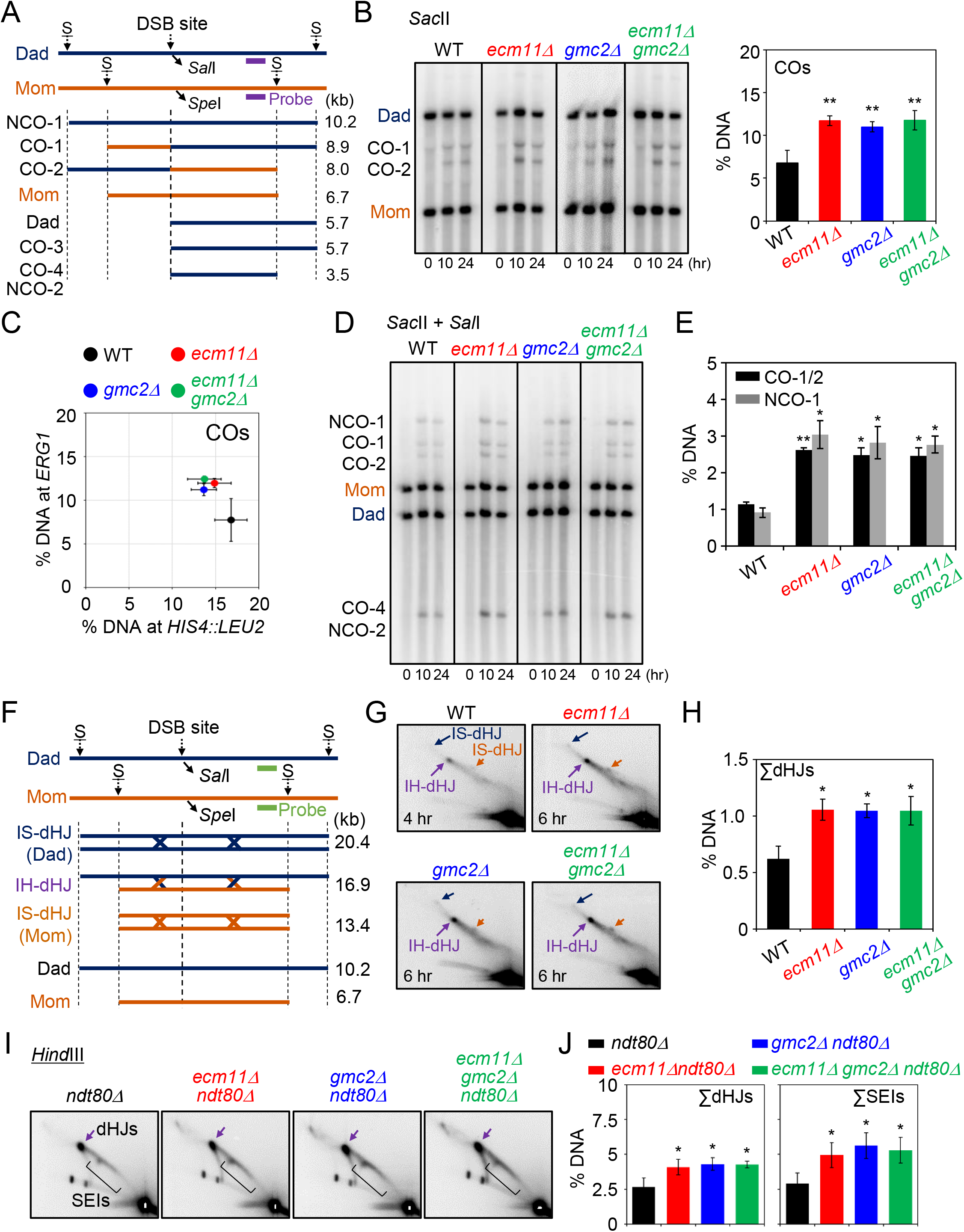
Meiotic recombination analysis at the *ERG1* locus in *ecm11Δ, gmc2Δ,* and *ecm11Δ gmc2Δ* mutants. (A) Schematic diagram of the *ERG1* locus showing restriction enzyme sites and position of the probe. Parental chromosomes, Mom and Dad, are distinguished by restriction enzyme site polymorphisms (S = *Sac*II). (B) Representative image of 1D gel analysis at the *ERG1* locus in WT, *ecm11Δ, gmc2Δ,* and *ecm11Δ gmc2Δ* strains. Quantitative analysis of the 1D gel at the *ERG1* locus in WT, *ecm11Δ, gmc2Δ,* and *ecm11Δ gmc2Δ* strains. CO levels are shown for maximum levels. Three independent meiotic cultures were used for calculation of the standard deviation (mean ± SD; *N* = 3). Significant differences were analyzed using unpaired *t*-tests (** *p* < 0.01). (C) Comparison of CO levels at the *HIS4LEU2* and *ERG1* loci. Each colored circle indicates the ratio of COs for *HIS4LEU2* versus *ERG1.* Data indicate mean ± SD (*N* = 3). (D) Representative image of CO and NCO analysis of WT, *ecm11Δ, gmc2Δ,* and *ecm11Δ gmc2Δ* strains. For CO and NCO gel analysis, the DNA samples were digested with *Sac*II and *Sal*I. (E) Quantitative analysis of CO and NCO. Data indicate mean ± SD (*N* = 3). Significant differences were analyzed using unpaired *t*-tests (\**p* < 0.05, \*\**p* < 0.01). (F) Physical map of the *ERG1* locus for 2D gel analysis. (G) Representative 2D gel analysis image of the *ERG1* locus. (H) Quantitative analysis of dHJs. Data indicate mean ± SD (*N* = 3). Significant differences were analyzed using unpaired *t*-tests (\**p* < 0.05). (I) Gel analysis (2D) of the *ERG1* locus in *ndt80Δ, ndt80Δ ecm11Δ, ndt80Δ gmc2Δ, and ndt80Δ ecm11Δ gmc2Δ* strains. The DNA samples were digested with *HindIII* restriction enzyme and used for 2D analysis to detect JMs at the *ERG1* locus. JMs in the *ERG1* locus were detected by Southern blotting using an *ERG1* probe (Lao et al., 2013). (J) Quantification of SEIs and dHJs at the *ERG1* locus. Data indicate mean ± SD (*N* = 3). Significant differences were analyzed by unpaired *t*-tests (\**p* < 0.05).

#### DSB frequencies

DSBs occurred on approximately 20% of chromatids at the *HIS4LEU2* locus, as estimated in a background strain (*rad50S*) where they failed to progress to form recombinants (Figure 2F; Supplementary Figure 4). In the *rad50S* background, the *ecm11Δ* and *gmc2Δ* mutants exhibited similar levels of DSBs at the *HIS4LEU2* locus. This was also confirmed in a *dmc1Δ* background where the DSB turnover was blocked (Supplementary Figure 5). This indicated that the mutations did not affect DSB formation at the *HIS4LEU2* locus during early meiosis.

#### Defects of SEI-dHJ transition and dHJ resolution

In all single and double *ecm11Δ gmc2Δ* mutants, DSBs formed at the *HIS4LEU2* locus with WT timing and eventually appeared to turnover similar to the WT strain (Figure 2C). However, CO formation in the mutants was delayed by approximately 2 h and CO levels reached only 80% of the WT levels, indicating a defect in JM processing to progress and form COs (Figure 2C). To confirm the JM-to-CO transition defects, we analyzed SEIs and dHJs using a native/native two-dimensional gel electrophoresis analysis followed by Southern hybridization. This revealed branched JMs of the recombination intermediates, in which IH JMs and intersister (IS) JMs could be distinguished (Figure 3A).

For the WT strain, SEIs and dHJs became detectable by 2D gel electrophoresis at 3.5 h and reached peak levels at 4 h with an IH:IS dHJ ratio of approximately 5:1. In the *ecm11Δ* and *gmc2Δ* mutants, SEIs and dHJs appeared at normal times and peaked at 6 h with a 2.5-h delay compared to those of the WT strain (Figure 3B; Supplementary Figure 6). Although SEIs and dHJs exhibited higher steady-state levels in *emc11Δ, gmc2Δ,* and *ecm11Δ gmc2Δ* cells between 5 h and 8 h relative to those in the WT, large portions of these JMs disappeared after 8 h (Figure 3B). However, unresolved SEI and dHJ species persisted in the *ecm11Δ* and *gmc2Δ* mutants at later times, which may have caused a defect in pachytene exit, and thus delayed the onset of meiosis I. The *ecm11Δ* and *gmc2Δ* mutants exhibited an IH:IS dHJ ratio of approximately 5.5:1, indicating normal IH bias in the mutants (Figure 3B). Overall, these results suggested that *emc11Δ, gmc2Δ,* and *ecm11Δ gmc2Δ* cells showed both a normal DSB-SEI transition and normal IH-bias, but had a defect in SEI-dHJ transition and/or dHJ resolution at the *HIS4LEU2* locus. Alternatively, these mutants may have formed more SEIs and dHJs, which could have resulted from more frequent DSB formation than what is observed in the WT strain.

To distinguish between these two possibilities, we examined the total number of dHJs in the *ecm11Δ* and *gmc2Δ* mutants in an *ndt80Δ* background, which causes meiotic cells to arrest in middle pachytene leading to the accumulation of SEIs and dHJs (Allers and Lichten 2001). The steady-state levels of dHJs at the *HIS4LEU2* locus in the *ecm11Δ ndt80Δ, gmc2Δ ndt80Δ,* and *ecm11Δ gmc2Δ ndt80Δ* mutants were similar to those in the *ndt80Δ* mutant (6.4 ± 0.8% in *ndt80Δ,* 6.2 ± 0.8% in *ndt80Δ ecm11Δ,* 6.5 ± 1.3% in *ndt80Δ gmc2Δ,* and 6.4 ± 1.0% in *ndt80Δ ecm11Δ gmc2Δ;* Figure 3C and 3D). This supported the hypothesis that the EG complex plays a positive role in SEI-dHJ transition and/or dHJ resolution, rather than in the regulation of JM frequencies.

#### *ecm11Δ* and *gmc2Δ* mutants demonstrate a locus-specific defect in DSB processing

As the effect of *ecm11Δ* and *gmc2Δ* mutations in meiotic recombination differed for chromosomes III and VII (Figure 1), we further analyzed meiotic recombination at the *ERG1* locus, which was identified as a natural hotspot in chromosome VII (Figure 4A). In contrast to the *HIS4LEU2* locus, the *ecm11Δ* and *gmc2Δ* mutants exhibited >2-fold increase in both CO and NCO at the *ERG1* locus (Figure 4A-4E). We then monitored JM formation at the *ERG1* locus using 2D gel electrophoresis and quantified the levels of JM species from parallel cultures of WT and *ecm11Δ* and *gmc2Δ* mutant cells (Figure 4F-4H; Supplementary Figure 7). The initiation time of JM formation in the mutants was similar to that in the WT strain, but the peak levels of SEIs and dHJs were approximately 3-fold higher in the *ecm11Δ* and *gmc2Δ* mutants (Figure 4H; Supplementary Figure 7). We then further analyzed JM formation at the *ERG1* locus in an *ndt80Δ* background. Interestingly, dHJ levels were increased from 2.65 ± 0.6% in *ndt80Δ* to 4.1 ± 0.5%, 4.3 ± 0.4%, and 4.3 ± 0.2% in the *ndt80Δ ecm11Δ, ndt80Δ gmc2Δ,* and triple mutants, respectively (Figure 4I and 4J). Therefore, in an *ndt80* background, the *ecm11Δ* and *gmc2Δ* mutants showed approximately 1.5-1.6-fold higher levels of dHJ relative to those in the control. This indicated an increased DSB event leading to JM formation. Thus, we interpreted these findings to signify that *ecm11Δ* and *gmc2Δ* mutants exhibited a combination of two defects in recombination at the *ERG1* locus. One defect resulted in increased JM formation (more establishment), while the other defect was in the SEI-dHJ transition and/or dHJ resolution (defective maintenance). The former defect was seen only for the *ERG1* locus but not for the *HIS4LEU2* locus. We further found that the *ecm11Δ* and *gmc2Δ* mutants showed slightly higher steady-state levels of DSBs at the *ERG1* locus relative to that of the WT (Supplementary Figure 8). In contrast, the levels of DSBs at the *ERG1* locus in the *ecm11Δ rad50S* and *gmc2Δ rad50S* mutants were similar to those in the *rad50S* mutant (Supplementary Figure 4).

### Early DSB formation is not affected by the absence of the EG complex

As noted above, elevated meiotic recombination in the *ecm11Δ* and *gmc2Δ* mutants may have been caused by increased initiation events associated with DSB formation. Using immunofluorescence analysis of chromosome spreads, we counted the number of foci of recombination proteins, such as Rad51 and Dmc1, as well as the number of ZMM foci such as Zip3 (Supplementary Figure 9). Foci formation by Rad51/Dmc1 and Zip3 in the *ecm11Δ* and *gmc2Δ* mutants began in a similar manner to that in the WT strain. However, the foci persisted longer on the chromosomes of the *ecm11Δ* and *gmc2Δ* mutants than on those of the WT strain, consistent with the delayed processing of recombination intermediates (Supplementary Figures 9).

We also checked the steady-state levels of DSBs at other loci, including *CYS3* (chromosome I), *ARG4* (chromosome VIII), and *BUD23* (chromosome III), in *rad5OS* and *dmc1Δ* backgrounds (Supplementary Figures 4 and 5). The *ecm11Δ* and *gmc2Δ* mutants showed similar DSB levels as the control strain at these three loci. These findings suggested that the EG complex does not play a role in DSB formation in early meiotic prophase I.

### EG complex restricts persistent DSB formation independent of Ndt80

It was previously reported that homolog engagement suppresses DSB formation in late prophase I (Thacker et al., 2014). In addition, pachytene exit mediated by Ndt80 also regulates DSB formation, which is independent of homolog engagement suppression (Thacker et al., 2014). We determined DSB levels in the absence of Ndt80 by quantifying Spo11-oligo complexes in a *gmc2Δ* background, which resulted in defective homolog engagement (Figure 5A and 5B). In WT cells, Spo11-oligos appeared and disappeared with a peak at 5 h. The *ndt80Δ* mutant exhibited persistent Spo11-oligos at late time points, consistent with the previous results (Thacker et al., 2014). Importantly, *gmc2Δ ndt80Δ* cells had increased levels of Spo11-oligos, with the increase being approximately 1.7-fold at 8 h compared to the levels in the *ndt80Δ* single mutant (Figures 5A and 5B). Consistent with this, Keeney and colleagues reported increased steady-state levels of Spo11-oligos in *gmc2Δ* and *ecm11Δ* mutants in a WT background, with the increased levels being more prominent at later times (Mu et al., 2020). These findings suggest that a greater degree of DSB formation occurs in late prophase I in *gmc2Δ*, which might be related to a homolog-engagement defect in the mutant. This phenomenon appears to be independent of Ndt80-mediated pachtyene exit. Sgs1 mutations promote chromosome synapsis in some synapsisdefective mutants called psuedosynapsis (Rockmill, 2003). Indeed, *sgs1-Δ200* mutation suppressed synapsis defects in *gmc2Δ* (Supplementary Figure 10). We evaluated whether pseudosynapsis could suppress DSB formation in the *gmc2Δ* mutant. However, Spo11-oligo complexes were increased approximately 1.7-fold in *sgs1-Δ200 gmc2Δ ndt80Δ* as seen in *gmc2Δ ndt80Δ* (Supplementary Figure 10). This suggested that the roles of the EG complex in suppression of DSB formation could not be replaced by pseudosynapsis.

**Figure 5.**
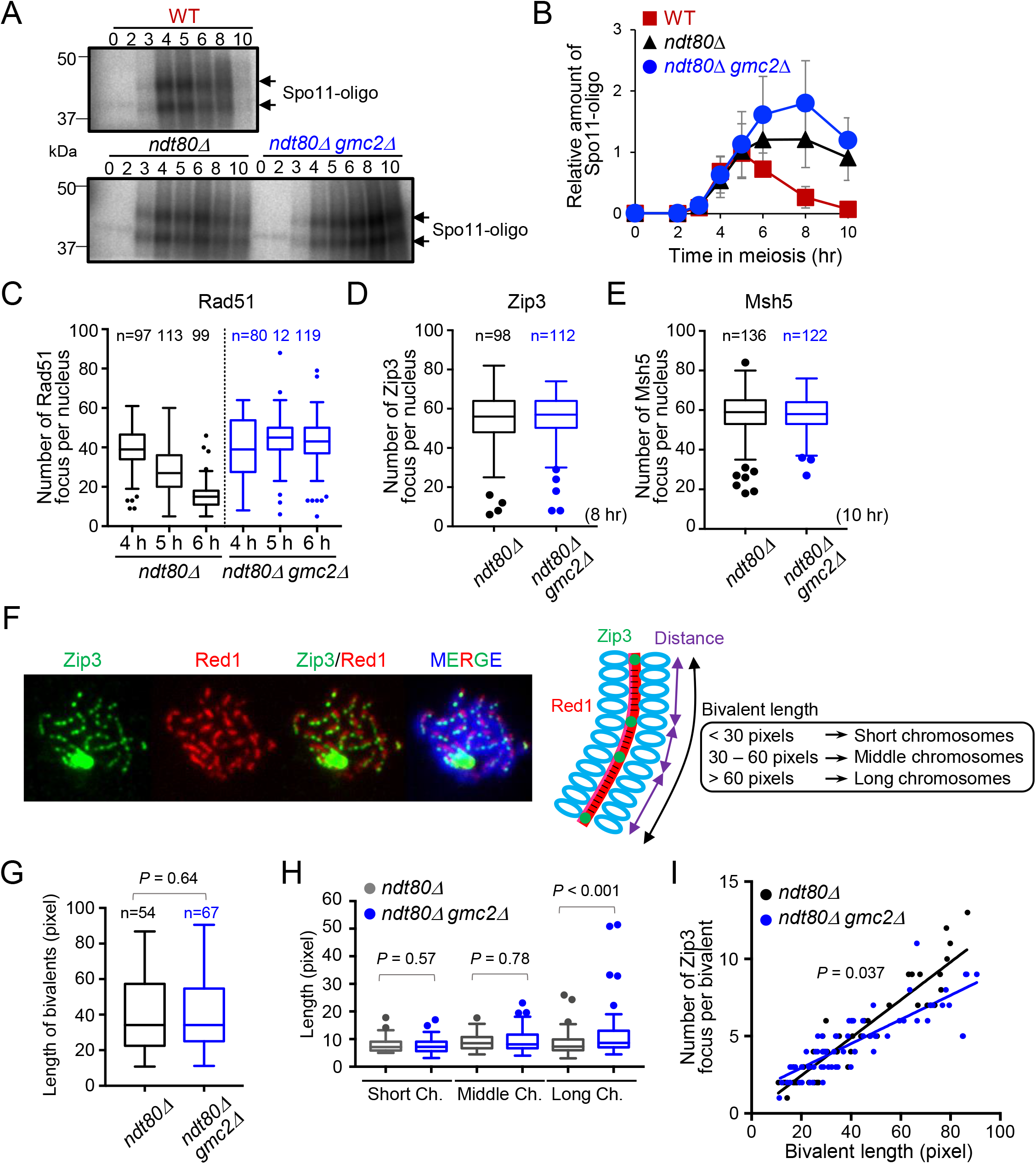
DSB formation and Zip3 distribution in WT and *gmc2Δ* cells. (A) Representative image of ^32^P-labeled DNA fragments covalently bound to Spo11-3FLAG in immunoprecipitates from WT, *ndt80Δ,* and *ndt80Δ gmc2Δ* cells at the indicated times. (B) Relative DNA fragment signals at each time point. Relative amounts of Spo11-oligo complex were calculated as described in Experimental Procedures. Data indicate mean ± SD (*N* = 3). (C) Average number of Rad51 foci per nucleus at the indicated time points analyzed for the *ndt80Δ* and *ndt80Δ gmc2Δ* mutants. The number of nuclei counted at each time point is shown at the top. (D) Average number of Zip3 foci per nucleus in the *ndt80Δ* and *ndt80Δ gmc2Δ* mutants. The number of nuclei counted at each timepoint is shown at the top. (E) Average number of Msh5 foci per nucleus in the *ndt80Δ* and *ndt80Δ gmc2Δ* mutants. The number of nuclei counted at each time point is shown at the top. (F) Representative image of meiotic nuclear spread from *ndt80Δ* cells at 8 hr post meiosis entry. The cells were co-stained for anti-Red1 (red) and anti-Zip3 (green). A schematic explanation of the classification of each category of bivalent length is shown. (G) Comparison of distribution of bivalent length between *ndt80Δ* and *ndt80Δ gmc2Δ* mutants. (H) Distribution of distances between adjacent Zip3 foci on short, medium, and long bivalents. Data indicate mean ± SD for more than three independent trials. The Mann□Whitney *U*-test was applied for statistical analysis and the results shown in panels G and H. (I) Correlations between the total numbers of Zip3 foci on each bivalent and the length of bivalent in WT and *gmc2Δ* strains. *P*-value was analyzed using Wald Wolfowitz runs test.

### EG complex regulates COs on long chromosomes

To determine the roles of the EG complex on pachytene chromosomes, we analyzed CO and synapsis formation at meiotic prophase I in the absence of Ndt80 and/or Gmc2 by immunostaining of chromosome spreads. The number of Rad51 foci was slightly increased and was maintained at late meiotic prophase in the *gmc2Δ ndt80Δ* double mutant compared with that in the *ndt80Δ* single mutant (Figure 5C). We also visualized Zip3 and Msh5 localization to detect CO formation in late prophase in the absence of Ndt80 and/or Gmc2. The number of Zip3 and Msh5 foci was similar in the *ndt80Δ* and *gmc2Δ ndt80Δ* mutants in late prophase (Figure 5D and 5E). However, more DSBs were produced in the mutants (Figure 5B). This implied that the *gmc2Δ* mutant produced additional DSBs in the late meiotic prophase, although these did not contribute to the total number of Zip3/Msh5-dependent recombinants in meiotic prophase I in an *ndt80Δ* background. It is likely that the additional DSBs in late meiosis of the mutant were repaired through a pathway that did not require Zip3/Msh5-focus formation.

To further explore the role of the EG complex in regulating CO control in a chromosome-dependent manner, bivalent length in an *ndt80Δ* background was revealed by staining for Red1, which localized to the chromosome axis at prophase, and was measured (Figure 5F). The total bivalent length indicated by Red1 lines in a single spread in *gmc2Δ ndt80Δ* was similar to that in *ndt80Δ (P =* 0.64; Figure 5G). This implied that normal axis formation occurred in the absence of Gmc2. We then measured the inter-distance of two adjacent Zip3 foci on a bivalent (Figure 5H) and counted the number of Zip3 foci per bivalent (Figure 5I). Bivalent length was classified for detectable Red1 signals according to chromosome length as follows: (1) short chromosomes (<30 pixels), (2) medium chromosomes (30–60 pixels), and (3) long chromosomes (>60 pixels) (Figures 5H and 5I). The short- and medium-length chromosomes displayed similar inter-distances between Zip3 foci and had a comparable number of foci per bivalent in the *ndt80Δ* and *gmc2Δ ndt80Δ* mutant strains (*P =* 0.57 and *P =* 0.78; respectively). Importantly, long chromosomes exhibited an increased inter-Zip3 distance in *gmc2Δ ndt80Δ* (more variation) compared with those in *ndt80Δ* (*P* < 0.001; Figure 5H). Furthermore, when the Zip3 foci number per bivalent was plotted against the chromosome length, the long chromosomes in *gmc2Δ ndt80Δ* showed a reduced Zip3 number compared with those in the control (Figure 5I). This suggested that ZMM-dependent events were less frequent on longer chromosomes in *gmc2Δ ndt80Δ* relative to that on other chromosomes. Therefore, we hypothesized that the high levels of COs on long chromosomes might have been caused by ZMM-independent recombination that originated in response to additional DSB formation, ultimately indirectly affecting the Zip3 foci number and distance.

### Absence of the EG complex suppresses the DSB turnover defect with *zip3* mutation

To further explore the EG complex in regulating CO formation, recombination intermediates of the *ecm11Δ* and *gmc2Δ* mutants, along their turnover, were determined in a *zip3Δ* background (Figure 6). In *zip3Δ* cells, DSBs remained at high levels at approximately 10–24 h at the *HIS4LEU2* locus (Figures 6A-6C), which was consistent with the findings of a previous report (Börner et al., 2004). Consistently, 2D gel analysis revealed that residual JMs still appeared at 24 h in *zip3Δ* (Figures 6D and 6E). By contrast, only low levels of DSBs were detected at approximately 10–24 h, and JMs were efficiently processed in *zip3Δ* cells without the EG complex (Figure 6C and 6E). Therefore, absence of the EG complex seemed to promote stalling of the DSB and/or JM processing in the *zip3Δ* mutant. Consistent with JM processing, COs accumulated to higher levels in all of the *ecm11Δ zip3Δ, gmc2Δ zip3Δ,* and *ecm11Δ gmc2Δ zip3Δ* mutants relative to those in the *zip3Δ* single mutant (Figure 6B). Consistent with the ability of double mutants to efficiently repair DSBs, meiotic divisions in the *ecm11Δ* and *gmc2Δ* mutants in a *zip3Δ* background occurred much earlier than those in the *zip3Δ* single mutant, implying that the absence of the EG complex partly ameliorated the defect in recombination progression caused by the absence of Zip3. Similar suppression of DSB-repair defects in the *zip3Δ* mutant by *ECM11* and/or *GMC2* deletion was observed at natural hotspots, including *ARG4, CYS3,* and *ERG1* loci (Supplementary Figure 11). Taken together, these results indicate that the EG complex could suppress recombination in the absence of Zip3, suggesting dual functions for the EG complex, i.e., promoting Zip3-dependent JM processing and suppressing Zip3-independent processing. Similar data were obtained when a *zip1* mutant was used instead of the *zip3* mutant (Supplementary Figure 12).

**Figure 6.**
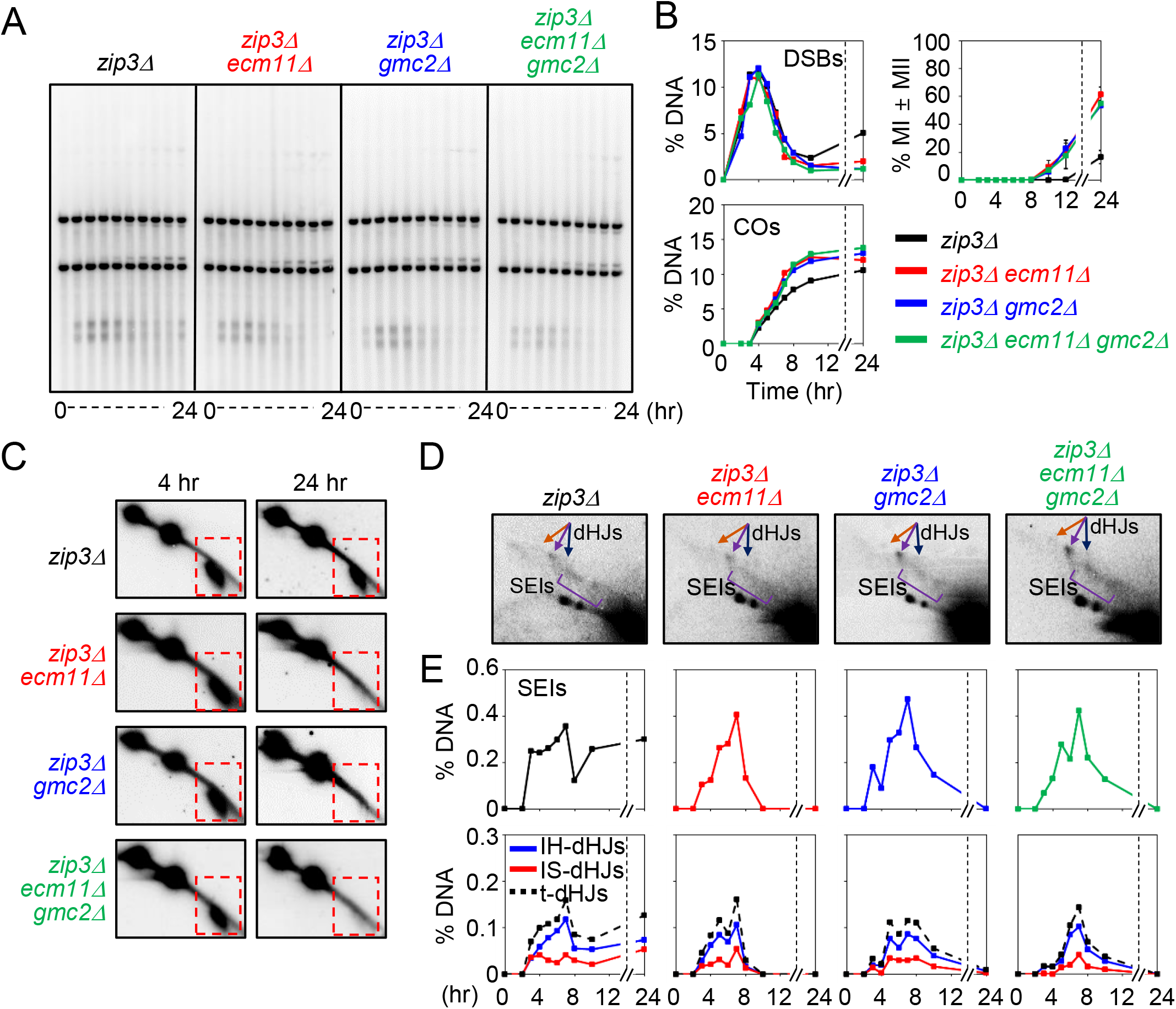
Ecm11 and Gmc2 inhibit additional DSB formation in *zip3Δ* cells. (A) Representative Southern blot image of 1D gel analysis in *zip3Δ, zip3Δ ecm11Δ, zip3Δ gmc2Δ,* and *zip3Δ ecm11Δ gmc2Δ* mutants assessed at 0, 2.5, 3.5, 4, 5, 6, 7, 8, 10, and 24 h. (B) Quantification of DSBs and total Cos, and analysis of meiotic division. (C) Two-dimensional gel detection of DSB formation at 4 h and 24 h in *zip3Δ, zip3Δ ecm11Δ, zip3Δ gmc2Δ,* and *zip3Δ ecm11Δ gmc2Δ* mutants. Dashed squares indicate DSB regions. (D) Representative image of 2D gel analysis in *zip3Δ, zip3Δ ecm11Δ, zip3Δ gmc2Δ,* and *zip3Δ ecm11Δ gmc2Δ* mutants. (E) Quantification of SEIs and dHJs from panel D.

### Suppression of recombination progression delays in *ecm11Δ* and *gmc2Δ* mutants

#### Low temperature alleviated progression delays

A previous study showed that high temperature introduces a kinetic block with respect to JM processing in the absence of ZMM, such as in case of *zip3Δ* (Börner et al., 2004). Therefore, we wondered whether temperature could affect the recombination defects in the *ecm11Δ* and *gmc2Δ* mutants, and evaluated the mutant phenotypes at a low temperature of 23°C (Supplementary Figure 13). Similar to the findings at 30°C, the *ecm11Δ* and *gmc2Δ* mutants exhibited reduced CO levels at the *HIS4LEU2* locus at 23°C without affecting the NCOs. In contrast, JMs did not accumulate at higher levels in the mutants compared to those in the WT strain at 23°C, and disappeared at later time points (Supplementary Figure 13). This indicated that low temperature suppressed the JM-processing defects in the *ecm11Δ* and *gmc2Δ* mutants. In other words, the kinetic barrier imposed by the EG complex is sensitive to temperature.

#### JM resolution under conditions of Cdc5 activation

It was previously shown that Cdc5, whose expression is induced by pachytene exit, promotes not only the disassembly of the SC and breakdown of SC proteins but also the resolution of the JMs to proceed to CO (Sourirajan and Lichten, 2008). The *ecm11Δ* and *gmc2Δ* mutants were defective in SC formation and in the JM-to-CO transition, which may induce checkpoint activation leading to suppressed *CDC5* expression. We hypothesized that ectopic expression of Cdc5 could induce an efficient resolution of CO intermediates in the *ecm11Δ* and *gmc2Δ* mutants. We evaluated recombination progression using specific conditional alleles in which the *CUP1* promoter was strongly activated in the presence of CuSO4 and replaced the normal promoter of *CDC5* (Supplementary Figure 14). When Cdc5 expression was induced at 6 h, JMs almost immediately began to disappear and there was an increase in the levels of COs (18% compared to 12% in the absence of CuSO_4_). This was consistent with the role of Cdc5 in JM resolution to COs. However, when Cdc5 was induced in the *ecm11Δ* and *gmc2Δ* mutants, which delayed JM progression, the JMs immediately resolved with a rapid increase in COs (Supplementary Figure 14). In contrast to that in the WT cells, forced Cdc5 expression did not increase the final level of COs in the mutants. These results suggested that in the absence of the EG complex, Cdc5 did not activate a canonical meiotic resolution of JMs to form COs. In the other words, the EG complex is critical for the biased resolution of JMs toward their progression to COs.

## DISCUSSION

The EG complex is a component of the SC central region and plays a role in its initiation and elongation. In the current study, we characterized *ecm11Δ* and *gmc2Δ* mutants, whose phenotypes provided new insights regarding the control of DSB formation along with ZMM-dependent and ZMM-independent recombination through homolog engagement.

### EG complex promotes the ZMM-dependent CO pathway

In normal meiosis, CO-designated DSBs are separated from NCO-fated DSBs during early meiosis prophase I (Börner et al., 2004; Kim et al., 2010). CO-designated DSBs are processed into JMs such as SEIs and dHJs. The dHJs are subsequently subjected to biased resolution into CO products. These JM-processing reactions are highly regulated in a meiosis-specific program and are coupled to morphological changes of chromosome structures, DSB-SEI, and SEI-dHJ transitions toward the resolution into COs, which are in turn roughly correlated with changes in SC morphology such as leptotene-zygotene, zygotene-early pachytene transition, and exit from the mid-pachytene, respectively (Börner et al., 2004; Hunter, 2006). Meiosis-specific ZMM proteins play a major role in JM processing into COs. In addition, there are mitotic processing pathways of JMs in meiotic cells, which are resolved into either COs or NCOs or are resolved into NCO (dissolution) (Dayani et al., 2011; De Muyt et al., 2012). A previous study further suggested that the “mitotic-like” Sgs1-dependent resolution of JMs is suppressed by ZMM proteins (De Muyt et al., 2012; Tang et al., 2016).

In the current study, physical analyses demonstrated that *ecm11Δ* and *gmc2Δ* mutants exhibited delayed JM processing such as in the SEI-dHJ transition and JM resolution. Interestingly, the DSB-SEI transition in these mutants appeared to be normal. This indicated that the EG complex was not required for early ZMM-dependent JM processing, but was needed for late processing, which might correlate with the establishment and maintenance of ZMM-dependent JM processing in meiosis (Figure 7). Consistent with that the *ecm11Δ* and *gmc2Δ* mutants showing normal establishment of the ZMM-pathway, cytological analysis in the present study showed normal Zip3 and Msh5 distributions on chromosomes in the *ecm11Δ* and *gmc2Δ* mutants. Conversely, ZMMs are required for the loading and polymerization of the EG complex together with the transverse filament protein Zip1. Taken together, the EG complex appears to be a positive modulator of late ZMM functions, particularly for the maintenance of ZMM-dependent recombination but not for its establishment. This is supported by the fact that *ecm11Δ* and *gmc2Δ* mutants did not affect NCOs, whose frequencies are indirectly affected by “early” ZMM functions (Börner et al., 2004).

**Figure 7.**
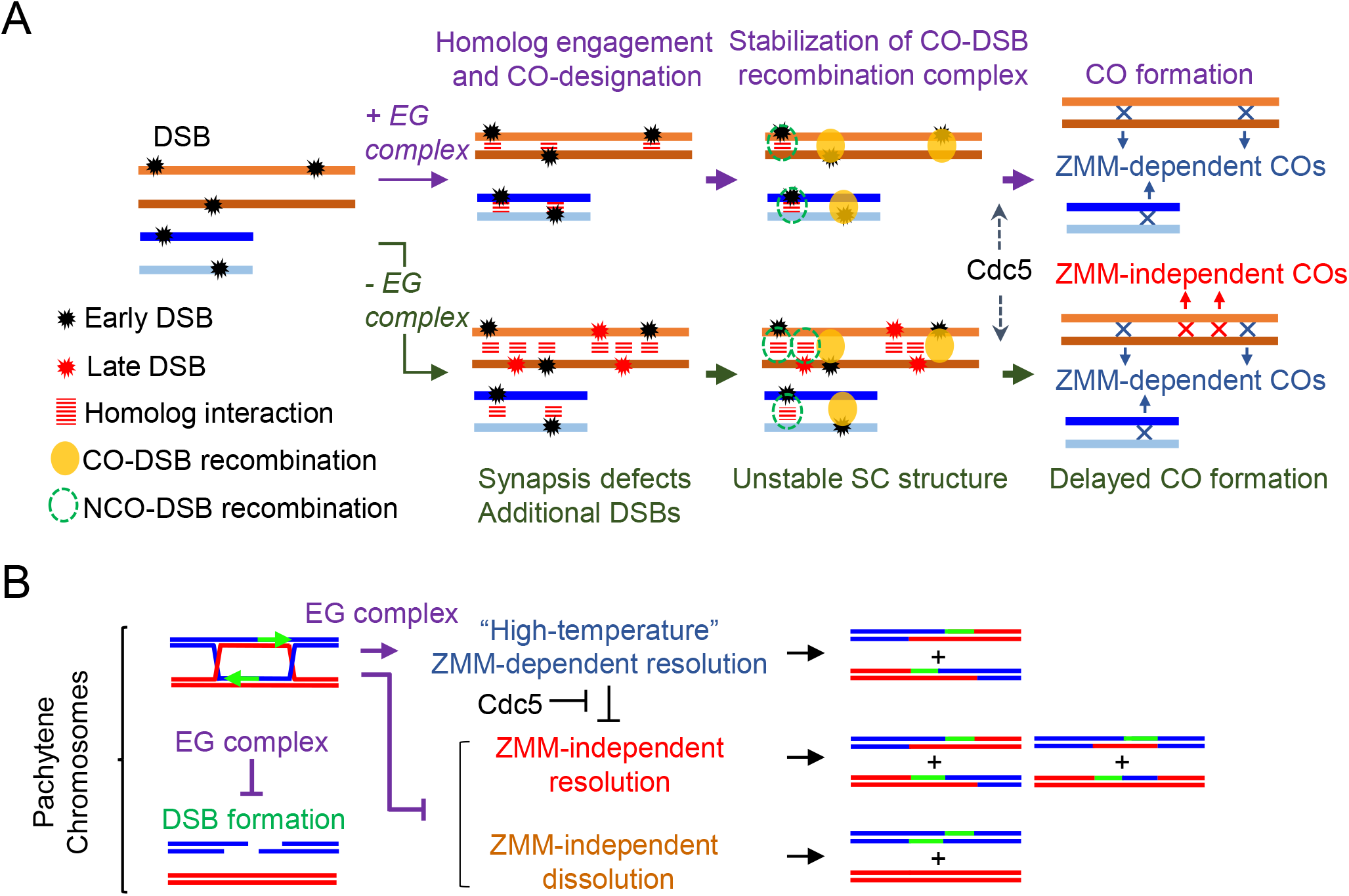
Roles of the EG complex in a feedback mechanism linked to DSB number and ZMM-dependent crossover formation. (A) Proposed mechanism of chromosome synapsis-dependent feedback as defined by *ecm11* and *gmc2* mutant phenotypes. EG complex facilitates the chromosomal assembly of Zip1 (Voelkel-Meiman et al., 2013) and modulates the meiotic recombination frequency and distribution through chromosome synapsis-dependent feedback. The arrest of JM resolution in the absence of the EG complex may be readily explained by a pathway in which unstable SC structures cause defection of CO-fated recombination. The EG complex-mediated SC central region provides an environment for proper recombination processing through phase separation. (B) Model for EG complex-mediated feedback controls of DSB formation, ZMM-dependent recombination, and ZMM-independent recombination.

### EG complex suppresses the ZMM-independent CO pathway

In the absence of the EG complex in a WT background, even with delayed processing of JMs, about two thirds of the JMs were resolved during late meiosis at approximately 7–8 h (Figure 2B). However, COs were gradually formed in the mutants. This suggested that a portion of the CO-designated JMs were not resolved into COs but rather into NCOs. This was consistent with the concept that the EG complex is required for the maintenance of ZMM functions. This resolution in the mutants may be independent of ZMM functions (Figure 7). In the absence of the EG complex, ZMM-independent processing of JMs seemed to operate for JM resolution, which could be catalyzed by mitotic resolvases. Indeed, we found that the *ecm11Δ* and *gmc2Δ* mutations suppressed a defect in JM processing in the *zip3Δ* and *zip1Δ* mutants. The *ecm11Δ zip3Δ* and *gmc2Δ zip3Δ* mutants formed more COs than the *zip3Δ* single mutant, but the amounts of COs in the double mutants were similar to those in the *ecm11Δ* and *gmc2Δ* single mutants. Given that the *ecm11Δ zip3Δ* and *gmc2Δ zip3Δ* mutants showed lower steady-state levels of JMs than the *ecm11Δ* and *gmc2Δ* mutants, most COs in the double mutants might not form through JM intermediates. This suggests that JM-processing activities of the probable “mitotic” resolvases seem to be more active in the absence of the EG complex than in its presence. In other words, the EG complex might limit the activity of “mitotic-like” JM processing enzymes not only in the absence of ZMM proteins but also in their presence.

### EG complex suppresses DSB formation in late prophase I

Genetic analysis showed increased CO and NCO frequencies on chromosome VII in the *gmc2Δ* mutant, which was consistent with previous genetic analyses of chromosome VIII (Voelkel-Meiman et al., 2016). This was further supported by physical analysis at the *ERG1* locus on chromosome VII as both CO and NCO levels were increased in the *ecm11Δ* and *gmc2Δ* mutants. These recombination increases may be simply be explained as being due to increased events of recombination initiation. Conversely, genetic analysis of markers on chromosome III showed decreased or WT-like levels of COs and slightly increased levels of NCO in the mutants. At the *HIS4LEU2* locus, reduced CO and normal NCO levels were observed in the *ecm11Δ* and *gmc2Δ* mutants. Furthermore, the formation of NCOs in the mutants was temporally separated from that of COs. Importantly, the levels of JMs at the locus in *ecm11Δ* and *gmc2Δ* mutants in an *ndt80Δ* background were the same as those in the parental *ndt80Δ* cells. Given that the mutants were defective in processing CO-designated DSBs, it was likely that at least “early” DSB levels on chromosome III in the mutants were the same or slightly increased relative to those in the WT strain. Indeed, when DSB levels were measured in repair-deficient mutants, such as *rad50S* and *dmc1Δ,* at least five loci on different chromosomes showed similar levels of DSBs between the WT, *ecm11Δ,* and *gmc2Δ* strains. This strongly suggested that the frequencies of early forming DSBs were not affected by absence of the EG complex. When the Spo11-oligo complex, which is a byproduct of DSB formation by Spo11 and whose amount is proportional to DSB frequencies, was analyzed in *ndt80Δ* and *gmc2Δ ndt80Δ* with a pachytene arrest, early amounts of Spo11-oligos and their kinetics were similar between the *ndt80Δ* and *gmc2Δ ndt80Δ* mutants. In contrast, the steady-state amount of Spo11-oligo at late meiotic prophase I increased more in the *gmc2Δ ndt80Δ* mutant than in *ndt80Δ.* This indicated that during late prophase I, more DSBs were formed in cells defective in the central region of the SC.

Previous studies have shown that more Spo11-oligos are observed in mutants defective in synapsis (e.g., *zmm* mutants), suggesting that homolog engagement suppresses DSB formation as a negative feedback control (Thacker et al., 2014; Kauppi et al., 2013). Our analysis of the *ecm11Δ* and *gmc2Δ* mutants supports this idea. In addition, our findings clearly indicated that ZMM assembly on meiotic chromosomes was not involved in this DSB suppression, as the *ecm11Δ* and *gmc2Δ* mutants showed normal Zip3/Msh5 foci formation. This indicated that SC elongation suppressed additional DSB formation in late prophase I as a feedback mechanism. In addition to the suppression by homolog engagement, DSB formation is negatively regulated by Ndt80 and recombination checkpoint kinases such as Tel1/ATM (Thacker et al., 2014; Zhang et al., 2011). Delayed JM processing in the *ecm11Δ* and *gmc2Δ* mutants may induce recombination checkpoints to downregulate Ndt80-dependent pachytene exit. As the Spo11-oligos levels were increased more in the *gmc2Δ ndt80Δ* mutant than in the *ndt80Δ* mutant, EG complex-dependent suppression of late DSBs apparently works independently of Ndt80. Activation of Tel1/ATM-dependent feedback control may explain the increased levels of DSBs in the *ecm11Δ* and *gmc2Δ* mutants. However, this is less likely as the levels of Spo11-oligos did not increase in the mutants during early meiosis when Tel1/ATM was activated. Moreover, in the background of the *rad50S* mutation, which robustly activates Tel1 kinase activity, we failed to observe any increase in DSB levels in the *ecm11Δ* and *gmc2Δ* mutants.

During late prophase I, SC elongation facilitates axis remodeling. The axis proteins Hop1 and Red1 are required for efficient DSB formation and are abundantly localized on chromosomes in a *zmm* mutant (Smith and Roeder, 1997). Furthermore, increased DSB levels were observed in the *ndt80Δ* mutant, which forms a full-length SC with Hop1/Red1 (Figure 5F) (Joshi et al., 2009). One possibility is that persistent Hop1/Red1 localization on chromosomes in the *ecm11Δ* and *gmc2Δ* mutants may induce additional DSBs. Therefore, EG complex-dependent suppression of DSB formation is likely to function through the removal of Hop1/Red1. In addition, the meiotic DSB-forming machinery might be functionally suppressed in the context of a full-length SC and the presence of the central regions.

### Role of SC central region in CO control

Synapsis-dependent suppression of DSB formation may explain increased recombination levels on long chromosomes. We speculated that the late-forming DSBs in unsynaptic chromosomes may be processed through a ZMM-independent recombination pathway that produces non-interfering COs and NCOs. The *ecm11Δ* and *gmc2Δ* mutants showed reduced CO interference relative to that of the WT in the genetic assays. In contrast, these mutants appeared to produce WT-like levels of Zip3 foci on chromosomes. Compromised CO interference in the mutants may be simply explained by the formation of additional non-interfering COs with adequate levels of interfering COs.

The fact that the *ecm11Δ* and *gmc2Δ* mutants retained significant CO interference with the normal number of Zip3 foci suggests that the establishment of CO interference is implemented in the absence of the central region of SC, and therefore in the absence of SC elongation or polymerization. Thus, SC polymerization and/or SC itself is not necessary for CO interference. This is consistent with the results presented by Kleckner and colleagues (Zhang et al. 2014a; Zhang et al. 2014b).

## Conclusion

Meiotic prophase I processes represent a unique meiotic event of the SC that mediates homologous chromosome pairing, homolog engagement, and crossing over via recombination (Börner et al., 2004; Storlazzi et al., 2010; Voelkel-Meiman et al. 2015). Little is currently known regarding the role of assembly of the SC central region in meiotic recombination. In this study, by analyzing the role of the EG complex in SC central region assembly, we determined that the EG complex-mediated SC central region was involved in multiple events pertaining to the control of recombination reactions, which ensured meiosis-specific properties such as regulated formation of interfering COs. The roles of the EG complex in controlling recombination include ZMM-dependent JM processing, JM resolution, and suppression of the ZMM-independent processing of JMs, as well as the downregulation of meiotic DSB formation during late prophase I (Figure 7). We propose that a compartment of the SC central region, which shows liquid-crystal properties and mediates phase separation (Rog et al., 2017), may function to sequester ZMM-dependent and ZMM-independent recombination proteins in the region, as well as to shuttle the DSB-forming machinery out of the region and that this is fostered by the EG complex and transverse filament Zip1.

## Experimental Procedures

### Yeast strains

All strains used in this study are derivatives of SK1. Strain genotypes are listed in Supplementary Table 1.

### Meiotic time courses

Meiotic time courses were studied as described previously (Hong et al., 2013; Oh et al., 2009; Kim et al., 2010; Yoon et al., 2016; Hong et al., 2019b). Strains maintained on YPG plates (1% yeast extract, 2% bactopeptone, 2% bactoagar, 3% glycerol) were streaked onto YPD plates (1% yeast extract, 2% bactopeptone, 2% bactoagar, 2% glucose) and grown for two days. A single colony was resuspended in 2 ml liquid YPD medium (1% yeast extract, 2% bactopeptone, 2% glucose) and grown to saturation. To induce synchronous meiosis, cells were inoculated into SPS medium (1% potassium acetate, 1% bactopeptone, 0.5% yeast extract, 0.17% yeast nitrogen base with ammonium sulfate and without amino acids, 0.5% ammonium sulfate, 0.05 M potassium biphthalate, pH 5.5) and incubated for 18 h. The cultures were then washed with pre-warmed SPM medium and resuspended in SPM medium (1% potassium acetate, 0.02% raffinose). Cells were harvested at indicated time points for each time course experiment. For the low-temperature time course experiments, synchronized cells were transferred to SPM medium and then the temperature was shifted to 23°C.

### Physical analysis

Genomic DNA was extracted from cultured cells using a guanidine-phenol extraction method as described previously (Kim et al., 2010; Hong et al., 2013; Yoon et al., 2016; Hong et al., 2019b). For physical analysis of JMs, cell cultures were harvested and cross-linked with psoralen under ultraviolet light. Genomic DNA (2 μg) was digested with 60 units *XhoI* restriction enzyme and electrophoresis was performed using a 0.6% agarose gel for 1D gel analysis. For native/native 2D gel analysis, 2.5 μg of *Xho*I-digested DNA samples was loaded onto 0.4% agarose gels, electrophoresed, and the gel lanes containing the DNA of interest were cut. The gel strips were then placed on 2D gel trays and 0.8% agarose containing ethidium bromide was poured into the trays. Two-dimensional gel electrophoresis was performed in Tris-borate-EDTA buffer containing ethidium bromide at approximately 6 V/cm for 6 h at 4°C. The gels were transferred to Biodyne B membranes (Pall) for Southern hybridization. The probes were radiolabeled with α-^32^P-dCTP using a random priming kit. Hybridizing DNA species were visualized using a phosphoimager (Bio-Rad) and quantified with QuantityOne software (Bio-Rad). For detection of the *HIS4LEU2* locus by Southern blotting, probes were amplified from yeast genomic DNA using primers 5’-ATATACCGGTGTTGGGCCTTT-3’ and 5’-ATATAGATCTCCTACAATATCAT-3’; primer sequences of DNA probes for the *ERG1* (*Sac*II, *Sac*II + *Sa/*I) locus were 5-ATGGAAGATATAGAAGGATACGAACC-3’ and 5’-GCGACGCAAATTCGCCGATGGTTTG-3’; and primer sequences of DNA probes for the *ERG1* (Hind*III*) locus were 5’-GGCAGCAACATATCTCAAGGCC-3’ and 5’-TCAATGTAGCCTGAGATTGTGGCG-3’.

### Spore viability and genetic distance

Spore viability and genetic distances between markers and CO interference were analyzed as previously described (Shinohara et al., 2008; Shinohara, 2019). Parental haploid strains (MSY4245 and MSY4304 derivatives) were mated for 3 h on YPAD plates and then transferred onto SPM plates. To exclude tetrads with mitotic COs, four independent crosses were analyzed. Map distances were calculated using the Perkins equation: cM = 100 (TT + 6NPD)/2(PD + TT + NPD). Standard errors were calculated using the Stahl Lab online tool (https://elizabethhousworth.com/StahlLabOnlineTools).

### Chromosome spreading and immunofluorescence

Immunostaining of yeast meiotic chromosome spreads was performed as described previously (Shinohara et al., 2000). Stained samples were observed using an epifluorescence microscope (Zeiss Axioskop 2) and a 100× objective (Zeiss AxioPlan, NA1.4). Images were captured using a CCD camera (Retiga; Qimaging) and processed using iVision (BioVision Technologies) and Photoshop (Adobe) software. Antibodies against Zip3 (rabbit and rat) (Shinohara et al, 2008), Rad51 (guinea pig) (Shinohara et al, 2000), Dmc1 (rabbit) (Hayase, 2004), Msh5 (rabbit) (Shinohara et al, 2008), and Red1 (chicken) (Shinohara et al, 2008) were generated using recombinant proteins purified from *Escherichia coli.*

### Spo11-oligo assay

Spo11-oligo detection was performed according to previously described methods (Neale and Keeney, 2009) with modifications. Spo11-oligo complexes were immunoprecipitated from 20 ml of synchronous meiotic yeast culture treated with 10% TCA. After preparation of the cell extract using glass beads (Yasui Kikai Co Ltd.), Spo11-FLAG was immunoprecipitated using anti-DYKDDDDK tag antibody (1E6, FUJIFILM Wako) and protein G-conjugated magnetic beads (Dynabeads, Veritas) in IP buffer (2% Triton X-100, 30 mM Tris-HCl [pH 8.0], 300 mM NaCl, 2 mM EDTA, 0.02% SDS). Immunoprecipitates were washed with IP buffer twice, and then a 10% volume of each sample was analyzed by western blotting and Spo11-FLAG protein levels in the precipitates were measured using an Odyssey infrared imaging system (LI-COR Biosciences). The remaining 90% of the samples were used for end-labeling reactions. For end-labeling of Spo11-oligo, immunoprecipitates were washed twice with NEBuffer #4 (New England Bio Labs). The beads were then suspended in TdT reaction buffer (1× NEBuffer #4, 0.25 mM CoCl2, 15 U TdT (Takara Bio), 20 Ci α-^32^P-dCTP [6000 Ci/mmol]), and incubated at 37°C for 2 h. Radio-labeled Spo11-oligos were separated by SDS-PAGE after washing with IP buffer three times, visualized using a Phosphor imager BAS5000 (FUJIFILM), and quantified using ImageQuant software (GE Healthcare).

## Acknowledgements

We are particularly thankful to Nancy Kleckner and Neil Hunter for providing the yeast strains. This work was supported by grants to K.P.K from the National Research Foundation of Korea funded by the Ministry of Science, ICT & Future Planning (No. 2020R1A2C2011887; 2018R1A5A1025077) and the Next-Generation BioGreen 21 Program (SSAC, No. PJ01322801), Rural Development Administration, Republic of Korea, and to M.S. from the Japan Society for the Promotion of Science (JSPS) KAKENHI (No. 19K22402 and 15H05973) and the Hyogo Science and Technology Association.

## Supplementary Information

**Supplementary Figure S1. CO interference analysis of WT and *gmc2Δ* strains**

(A) CO interference in NPD ratio in chromosomes III, VII and V.

(B) Coefficient of coincidence in chromosomes III and VII.

**Supplementary Figure S2. Spore viability test for WT, *ecm11Δ, gmc2Δ* and *ecm11Δ gmc2Δ* strains**

(A) Meiotic nuclear division for WT, *ecm11Δ, gmc2Δ* and *ecm11Δ gmc2Δ* strains.

(B) Spore viability analysis for WT, *ecm11Δ, gmc2Δ* and *ecm11Δ gmc2Δ* strains.

**Supplementary Figure S3. Gel analysis (2D) of CO and NCO for *ecm11Δ* and *gmc2Δ* mutants**

(A) Representative image of two-dimensional (2D) gel analysis of CO and NCO. Genomic DNA was digested with *Xho*I restriction enzyme for first dimension gel analysis and digested *in situ* with *BamHI* for second dimension gel analysis.

(B) Quantitative analysis of the 2D gel of CO and NCO in WT, *ecm11Δ, gmc2Δ* and *ecm11Δ gmc2Δ* strains.

**Supplementary Figure S4. Analysis of DSB levels in *rad50s* backgrounds**

(A) Images (1D) of the *HIS4LEU2* locus in *rad50S, rad50S ecm11Δ, rad50S gmc2Δ* and *rad50S ecm11Δ gmc2Δ* cells (left). Quantification of DSBs from three independent meiotic cultures (right).

(B) Gel analysis (1D) at different loci in *rad50S, rad50S/ecm11Δ, rad50S gmc2Δ* and *rad50S ecm11Δ gmc2Δ* strains. *ARG4, BUD23, CYS3,* and *ERG1* loci located on chromosomes VIII, III, I and VII, respectively.

(C) Quantitative analysis of DSBs at various loci in three and two (*ERG1)* sets of independent meiotic cultures.

**Supplementary Figure S5. Analysis of DSB levels in *dmc1Δ* backgrounds**

(A) Gel analysis (1D) at the *HIS4LEU2, ARG4, BUD23,* and *CYS3* loci in *dmc1Δ, dmc1Δ ecm11Δ, dmc1Δ gmc2Δ* and *dmc1Δ ecm11Δ gmc2Δ* mutants.

(B) Quantification of DSBs.

**Supplementary Figure S6. Gel analysis (2D) for WT, *ecm11Δ, gmc2Δ,* and *ecm11Δ gmc2Δ* strains at the *HIS4LEU2* hotspot**

(A) Gel images (2D) of Southern blotting for WT, *ecm11Δ, gmc2Δ* and *ecm11Δ gmc2Δ* strains at the *HIS4LEU2* locus. Images show DNA species from representative meiotic time courses. (B) Representative images and quantitative analysis of WT, *ecm11Δ, gmc2Δ* and *ecm11Δ gmc2Δ* strains at the *HIS4lEU2* locus.

**Supplementary Figure S7. Gel analysis (2D) for WT, *ecm11Δ, gmc2Δ* and *ecm11Δ gmc2Δ* strains at the *ERG1* locus**

(A) Gel images (2D) of Southern blotting for WT, *ecm11Δ, gmc2Δ* and *ecm11Δ gmc2Δ* strains at the *ERG1* locus. Images show DNA species from representative meiotic time courses.

(B) Representative images and quantitative analysis of WT, *ecm11Δ, gmc2Δ* and *ecm11Δ gmc2Δ* strains at the *ERG1* locus.

**Supplementary Figure S8. Analysis of DSB levels in WT, *ecm11Δ, gmc2Δ* and *ecm11Δ gmc2Δ* at the *ERG1* locus**

(A) Representative 1D gel images of WT, *ecm11Δ, gmc2Δ* and *ecm11Δ gmc2Δ* at the *ERG1* locus.

(B) Quantitative analysis of images shown in (A). Error bars indicate mean ± SD (N = 2).

**Supplementary Figure S9. Chromosome analysis of WT, *ecm11Δ, gmc2Δ* and *ecm11Δ gmc2Δ* cells**

(A) Representative images of chromosome spreads and cells immunostained for Zip3 (green) along meiotic progression of WT and *gmc2Δ* cells.

(B) Quantification of the number of Zip3 foci-positive nuclei along meiotic progression in WT (black), *ecm11Δ* (red), *gmc2Δ* (blue) and *ecm11Δ gmc2Δ* (green) cells.

(C) Quantification of the number of Zip3 foci along meiotic progression in WT and *gmc2Δ* cells.

(D) Representative images of chromosome spreads and cells immunostained for Rad51 (green) and Dmc1 (red) along meiotic progression in WT and mutant cells.

(E) Quantification of the number of Rad51 and Dmc1 foci-positive nuclei along meiotic progression.

(F) Quantification of the number of Rad51 and Dmc1 foci along meiotic progression.

**Supplementary Figure S10. EG complex prevents additional DSB formation after the Ndt80 pathway, even with pseudosynapsis**

(A) Representative image of meiotic nuclear spread from *ndt80Δ gmc2Δ* and *ndt80Δ gmc2Δ sgs1-Δ200* cells at 8 hr post meiosis entry. Cells were co-stained for anti-Red1 (green), antiZip1 (red), and DAPI (blue). Schematic presentation of chromosome structures of each mutant.

(B) Representative image of ^32^P-labeled DNA fragments covalently bound to Spo11-3FLAG in immunoprecipitates and quantitative analysis of the images for *ndt80Δ*, *ndt80Δ gmc2Δ*, *ndt80Δ sgs1-Δ200* and *ndt80Δ gmc2Δ sgs1-Δ200* mutants. Error bars indicate mean ± SD (*N* = 4).

**Supplementary Figure S11. Absence of EG complex restrains *zip3Δ*-induced additional DSBs at various loci**

(A, B, and C) Southern blot analysis (1D) of *ARG4, CYS3,* and *ERG1* loci in *zip3Δ, zip3Δ ecm11Δ, zip3Δ gmc2Δ* and *zip3Δ ecm11Δ gmc2Δ* mutants.

(D) Quantitative analysis of DSB shown in panels A, B and C.

**Supplementary Figure S12. Absence of EG complex restrains *zip3Δ*-induced additional DSBs at various loci**

(A, B, and C) Southern blot analysis of *ARG4, CYS3,* and *ERG1* loci in *zip1Δ, zip1Δ ecm11Δ, zip1Δ gmc2Δ* and *zip1Δ ecm11Δ gmc2Δ* mutants.

(D) Quantitative analysis of DSB shown in panels A, B and C.

**Supplementary Figure S13. Meiotic recombination of WT, *ecm11Δ, gmc2Δ* and *ecm11Δ/gmc2Δ* mutants at low temperature**

(A) CO/NCO analysis of WT, *ecm11Δ, gmc2Δ* and *ecm11Δ gmc2Δ* strains at low temperature.

(B) Representative images of 2D gel Southern blotting time course for WT, *ecm11Δ, gmc2Δ* and *ecm11Δ gmc2Δ* strains at 23°C.

(C) Quantitative analysis shown in panel B

**Supplementary Figure S14. Expression of Cdc5 ameliorates pachytene arrest in *ecm11Δ* or *gmc2Δ* mutants**

(A) Representative images of 1D gel for *P_CUP1_-CDC5, P_CUP1_-CDC5 ecm11Δ* and *P_CUP1_-CDC5 gmc2Δ* in the absence and presence of CuSO_4_. A total of 30 μM CuSO_4_ was added to each meiotic culture at 6 hr post induction of meiosis.

(B) Quantification of COs.

(C) Representative image of 2D gel analysis of *P_CUP1_-CDC5, P_CUP1_-CDC5 ecm11Δ* and *P_CUP1_-CDC5 gmc2Δ* in the presence or absence of CuSO_4_.

(D) Quantification of SEIs and dHJs. Arrows indicate the time for inducing Cdc5 expression.

**Supplementary Table 1.**
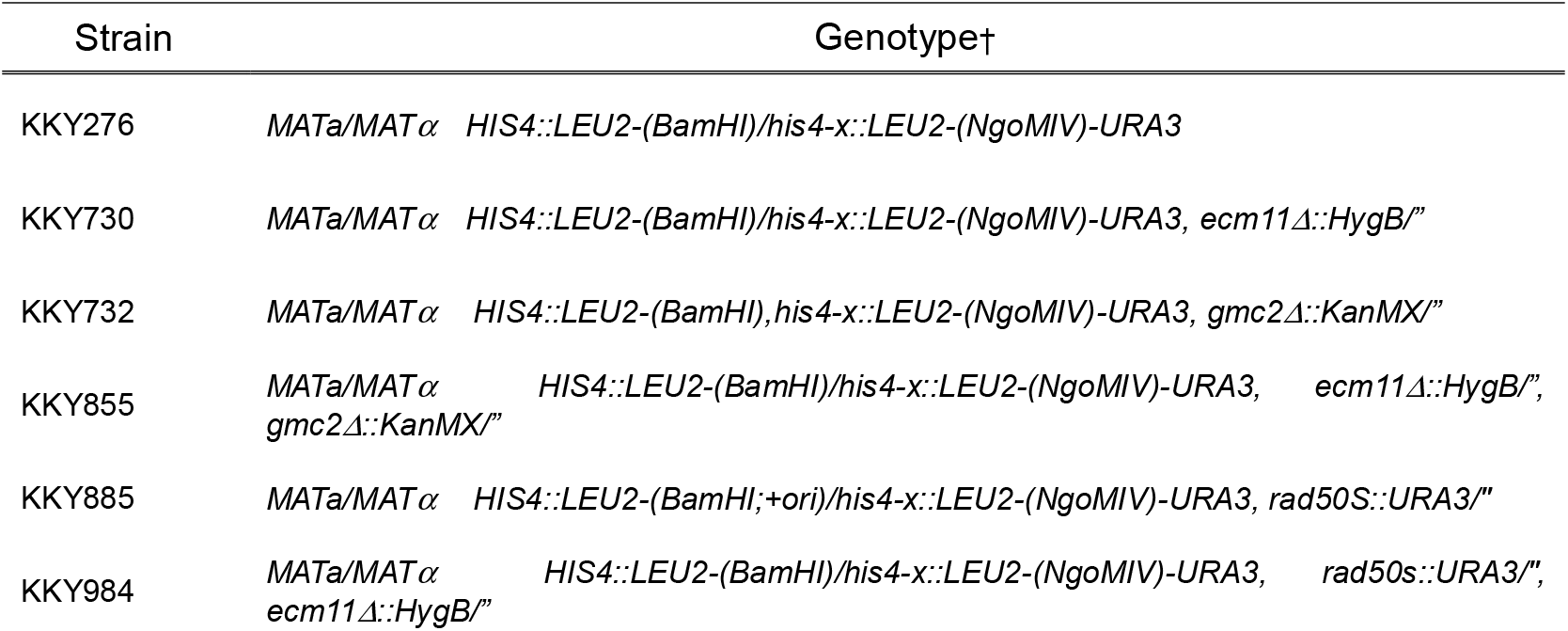
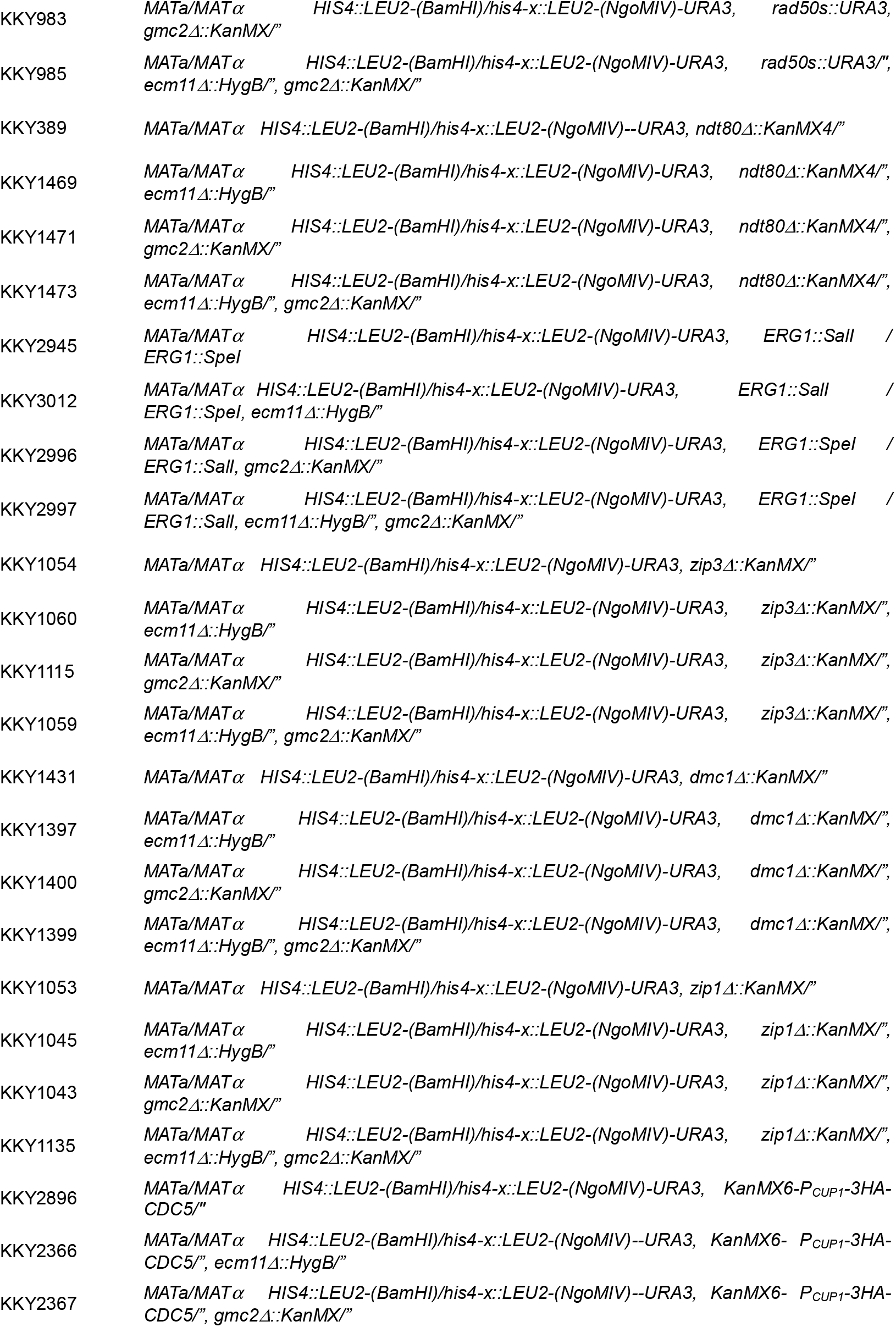
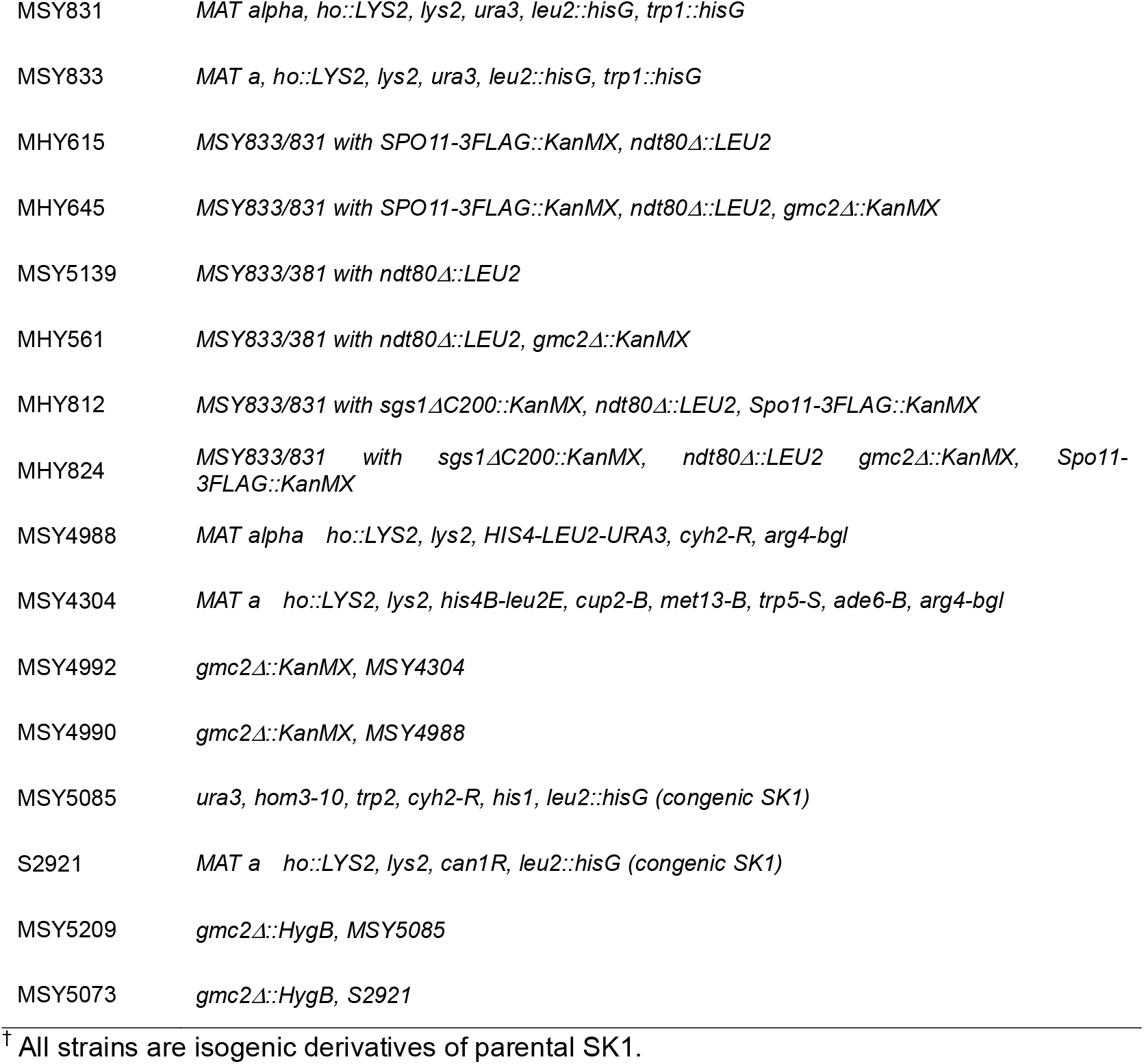
Yeast strains used in this study.

