## Supplementary figures for "The Synaptonemal Complex Central Region Modulates Crossover Pathways and Feedback Control of Meiotic Double-strand Break Formation"

### Supplemental Figure 1

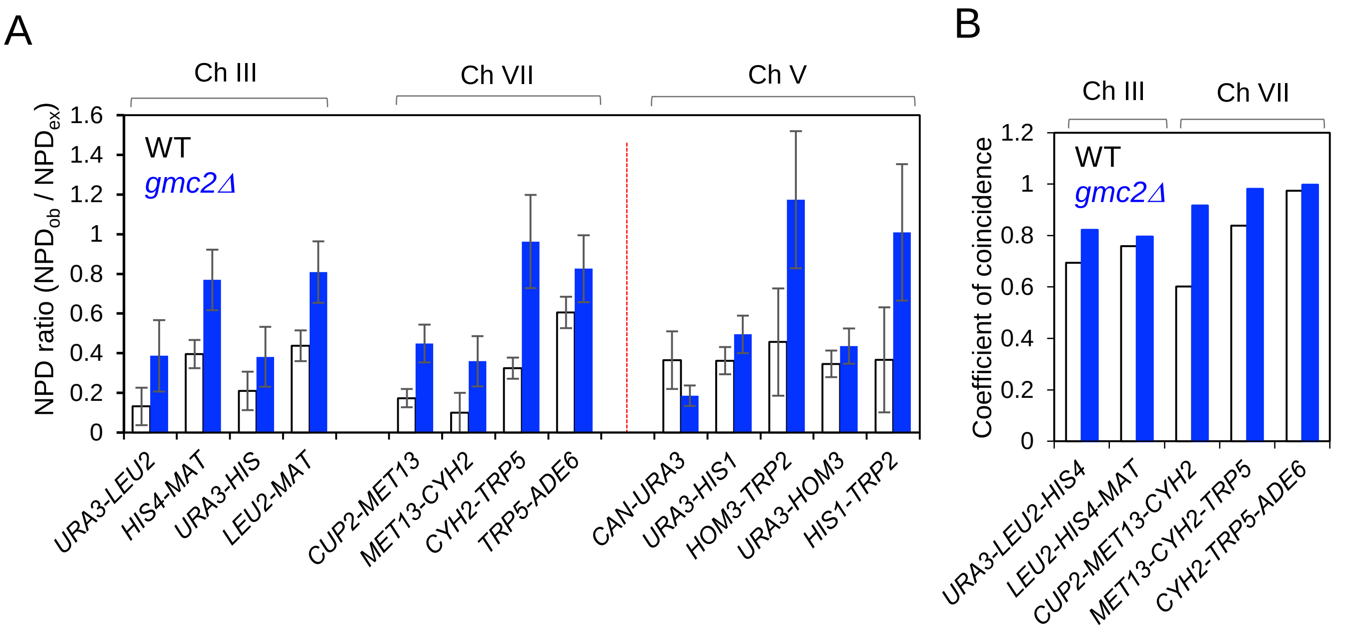

Supplemental Figure 2

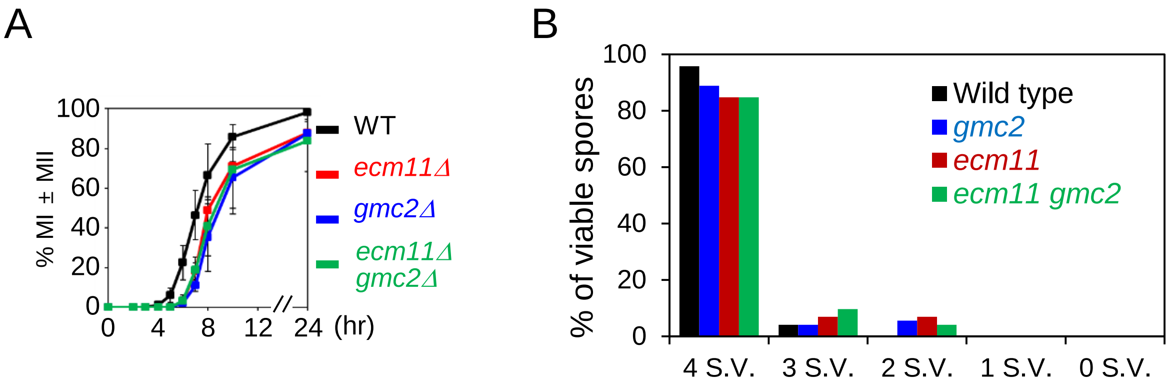

### Supplemental Figure 3

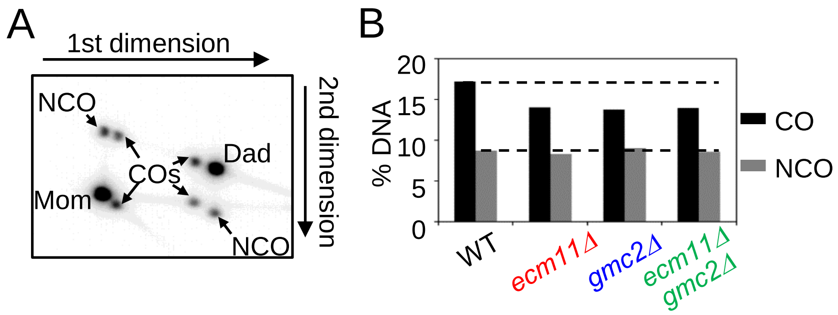

### Supplemental Figure 4

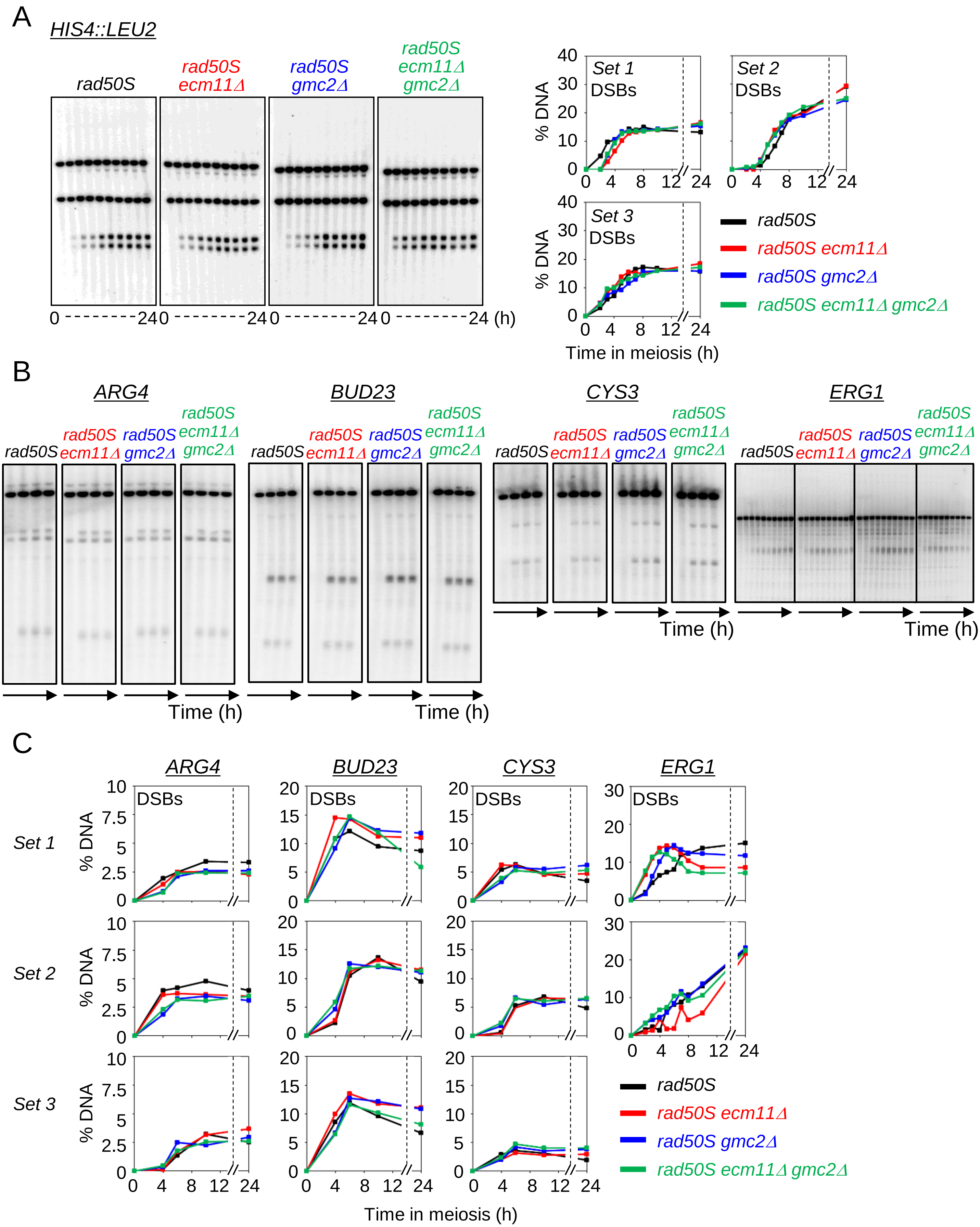

Supplemental Figure 5

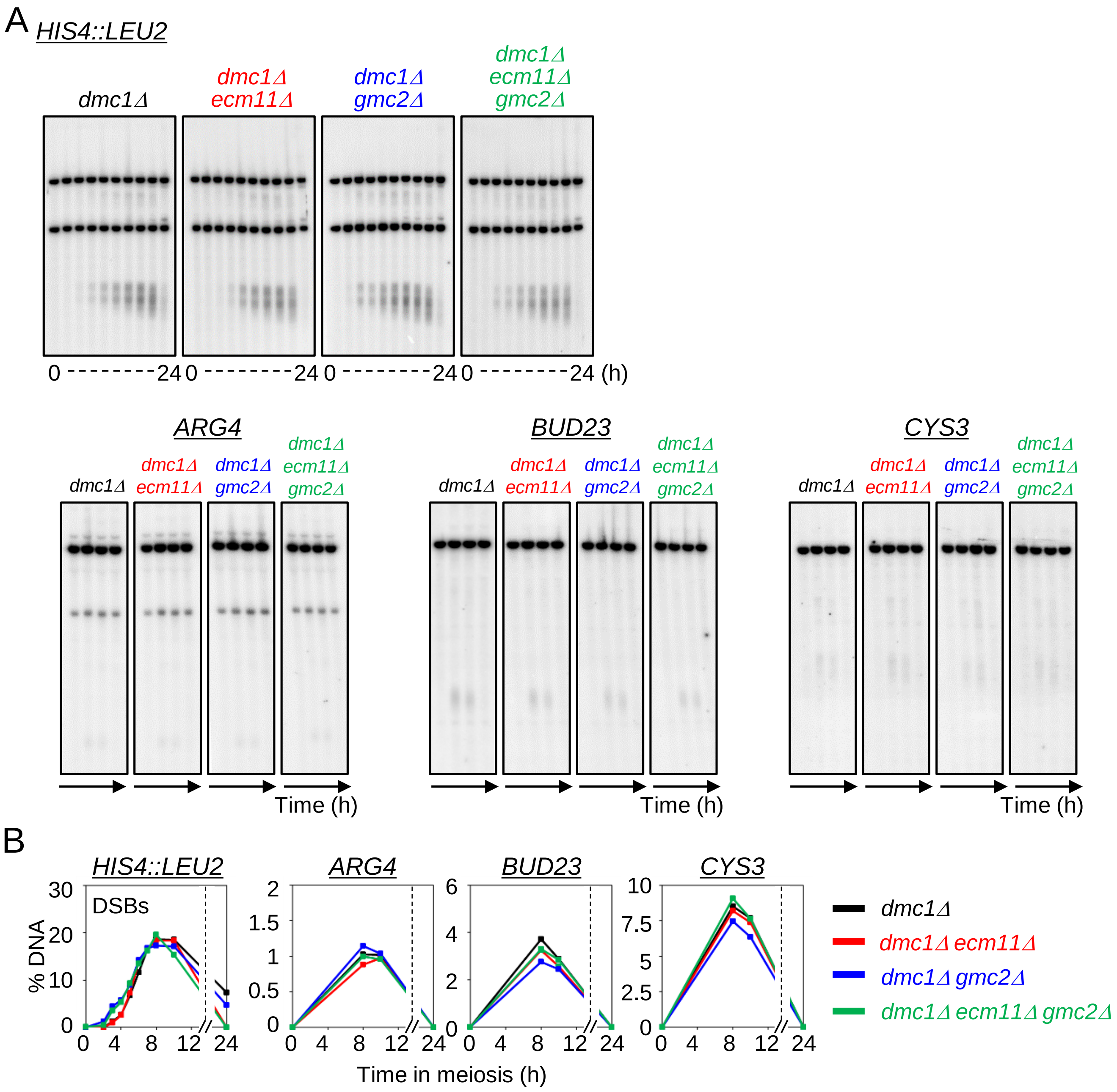

### Supplemental Figure 6

A

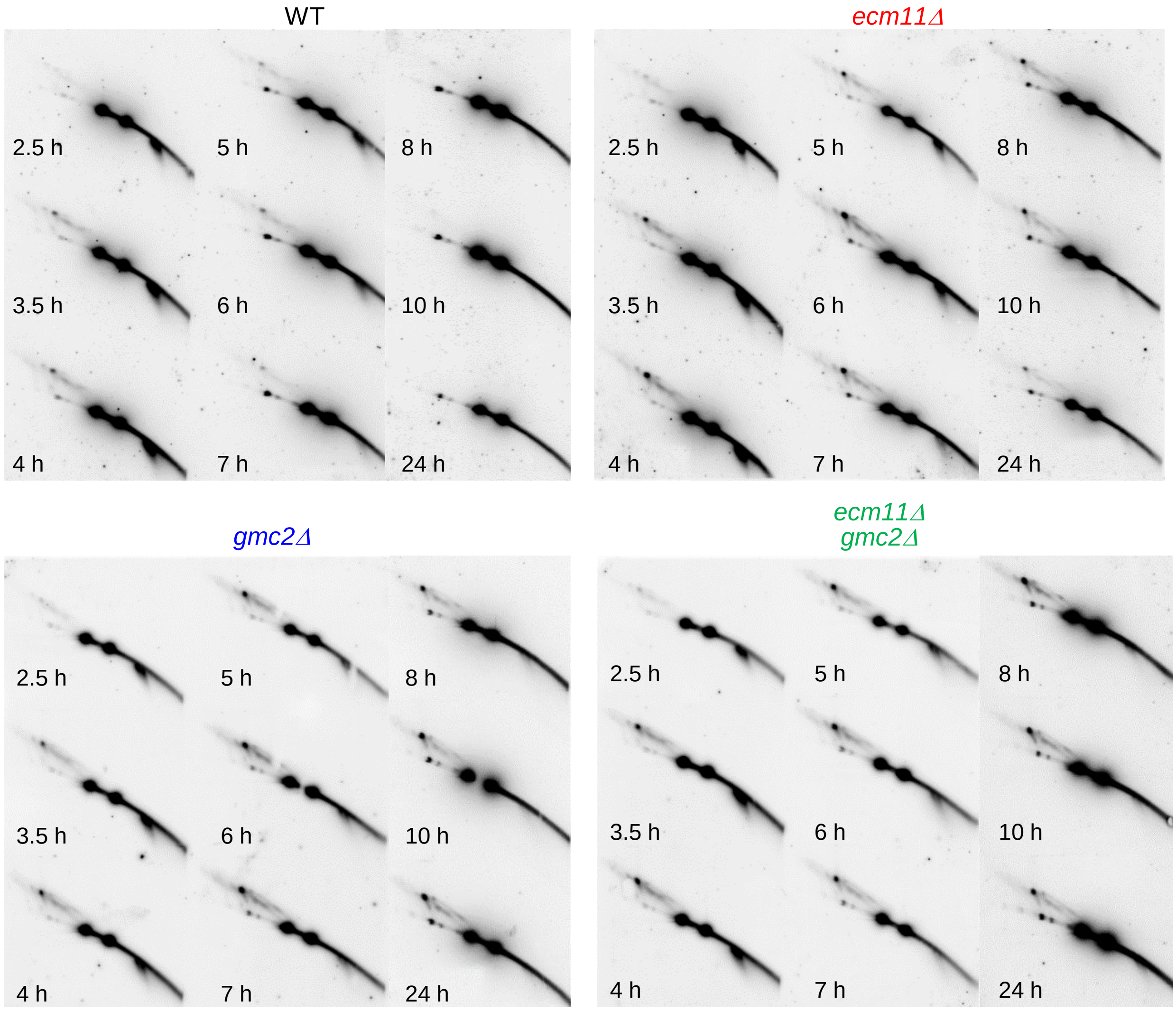


B

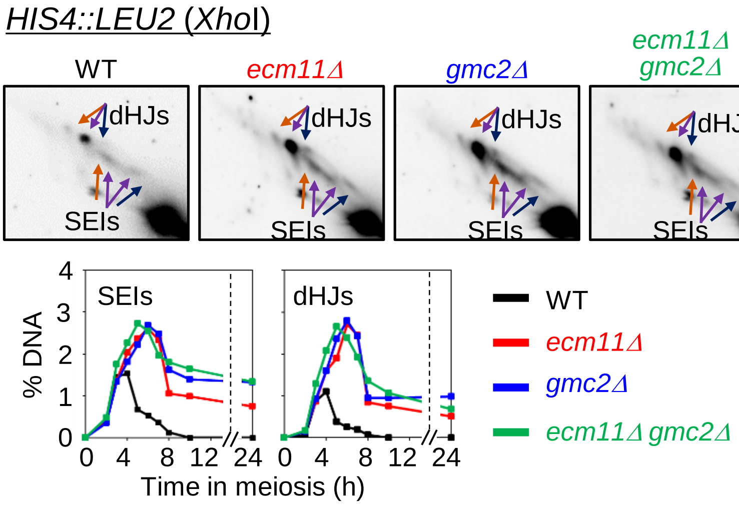

### Supplemental Figure 7

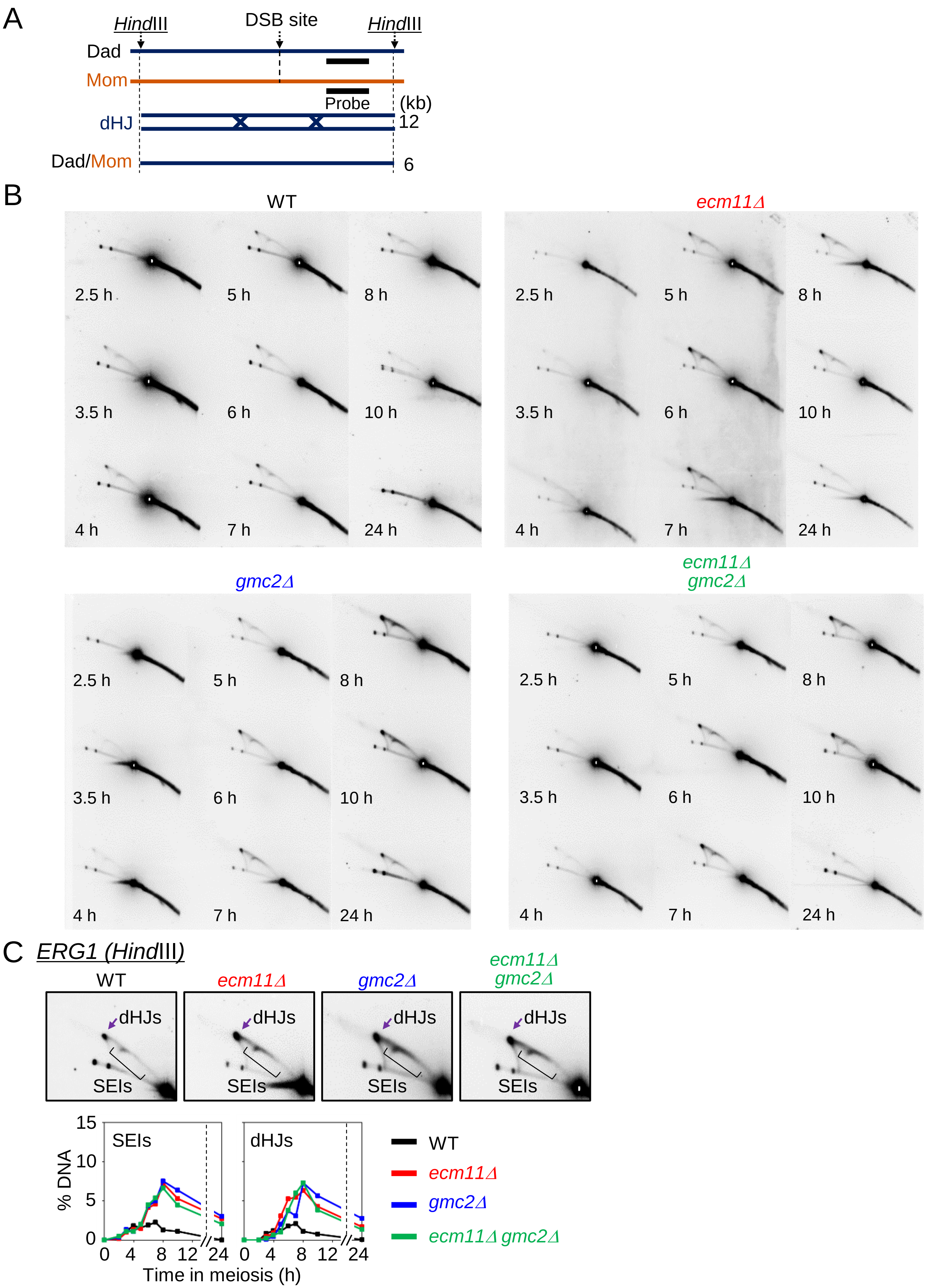

### Supplemental Figure 8

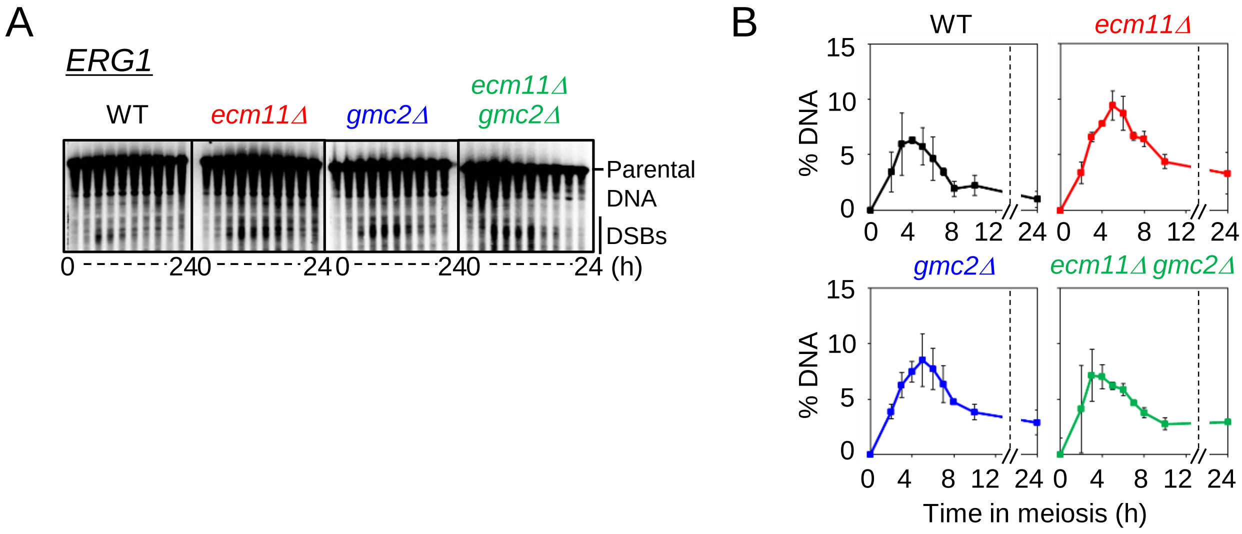

### Supplemental Figure 9

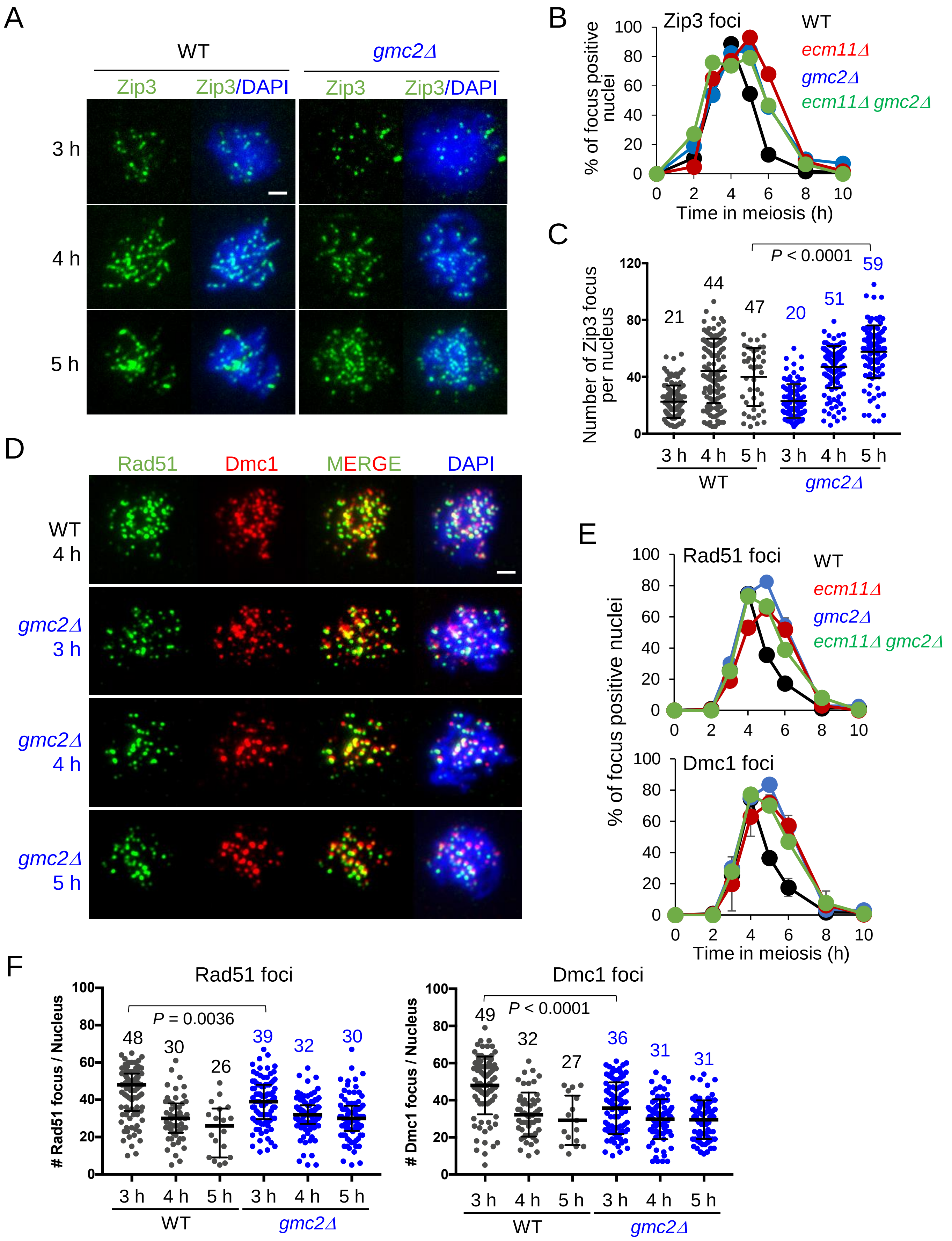

Supplemental Figure 10

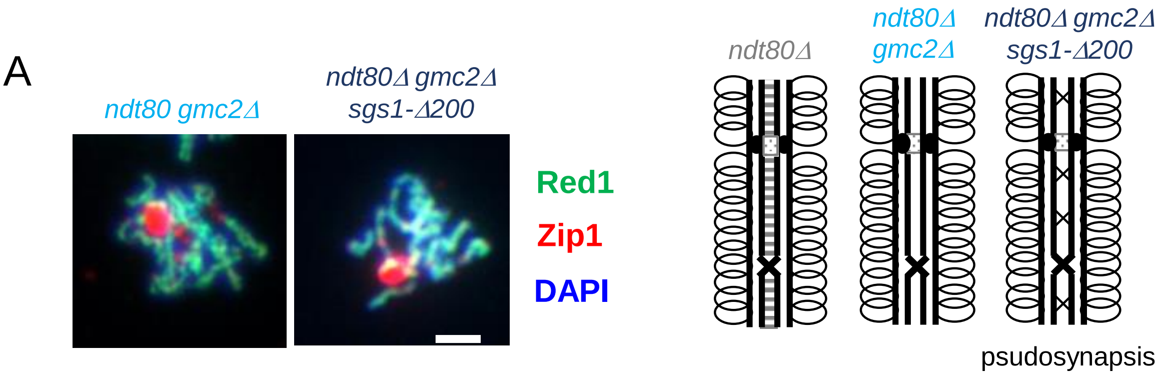


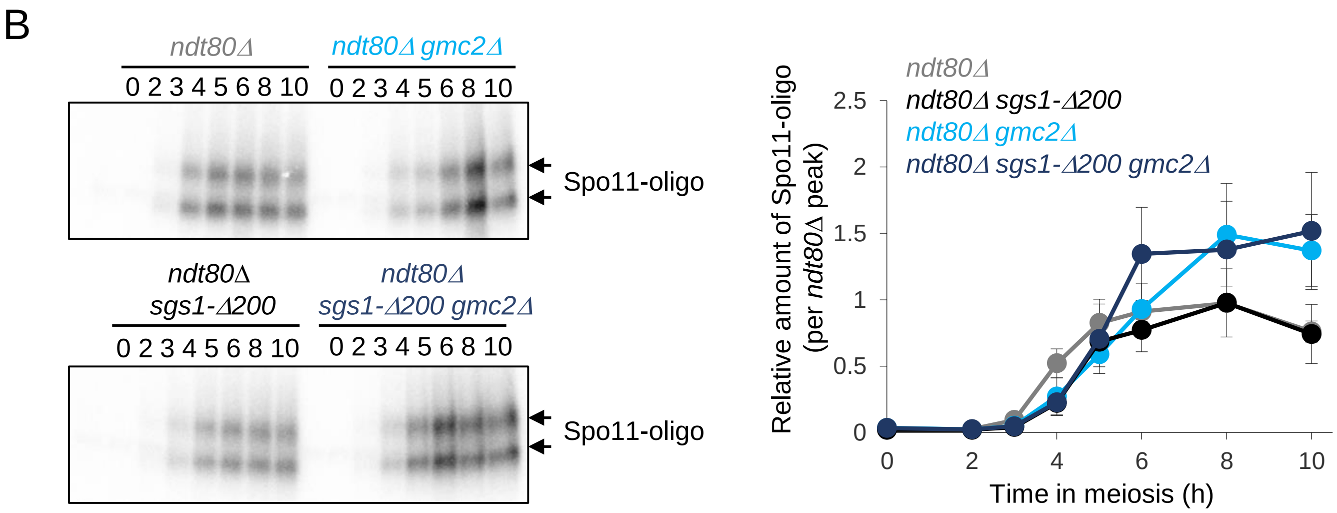

### Supplemental Figure 11

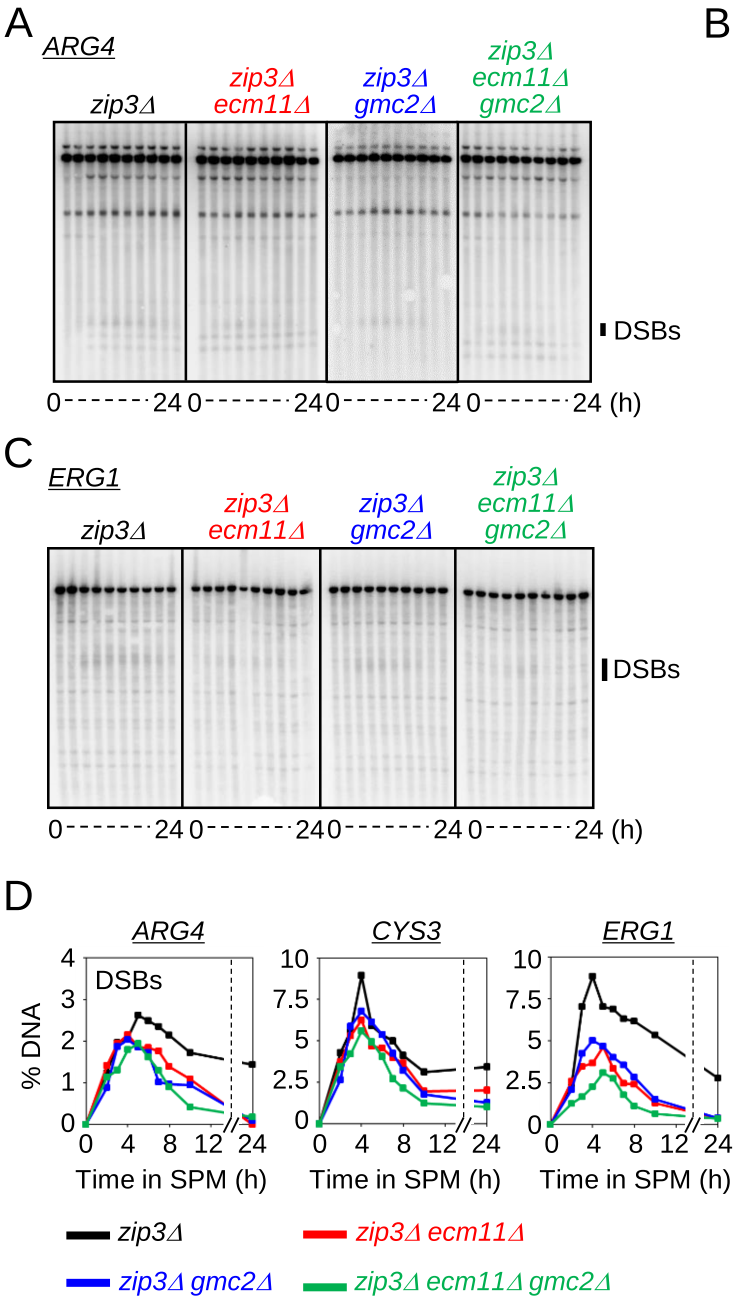

### Supplemental Figure 12

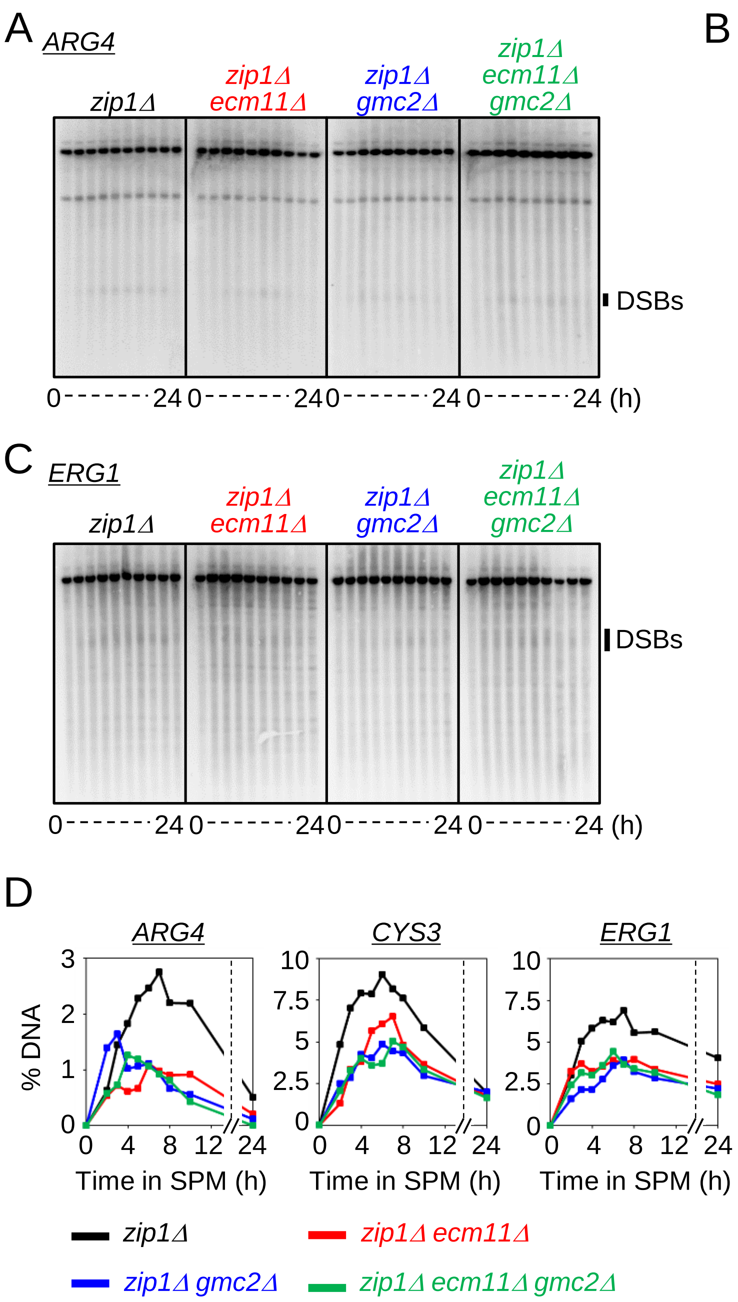

### Supplemental Figure 13

#### 23ºC meiotic time course

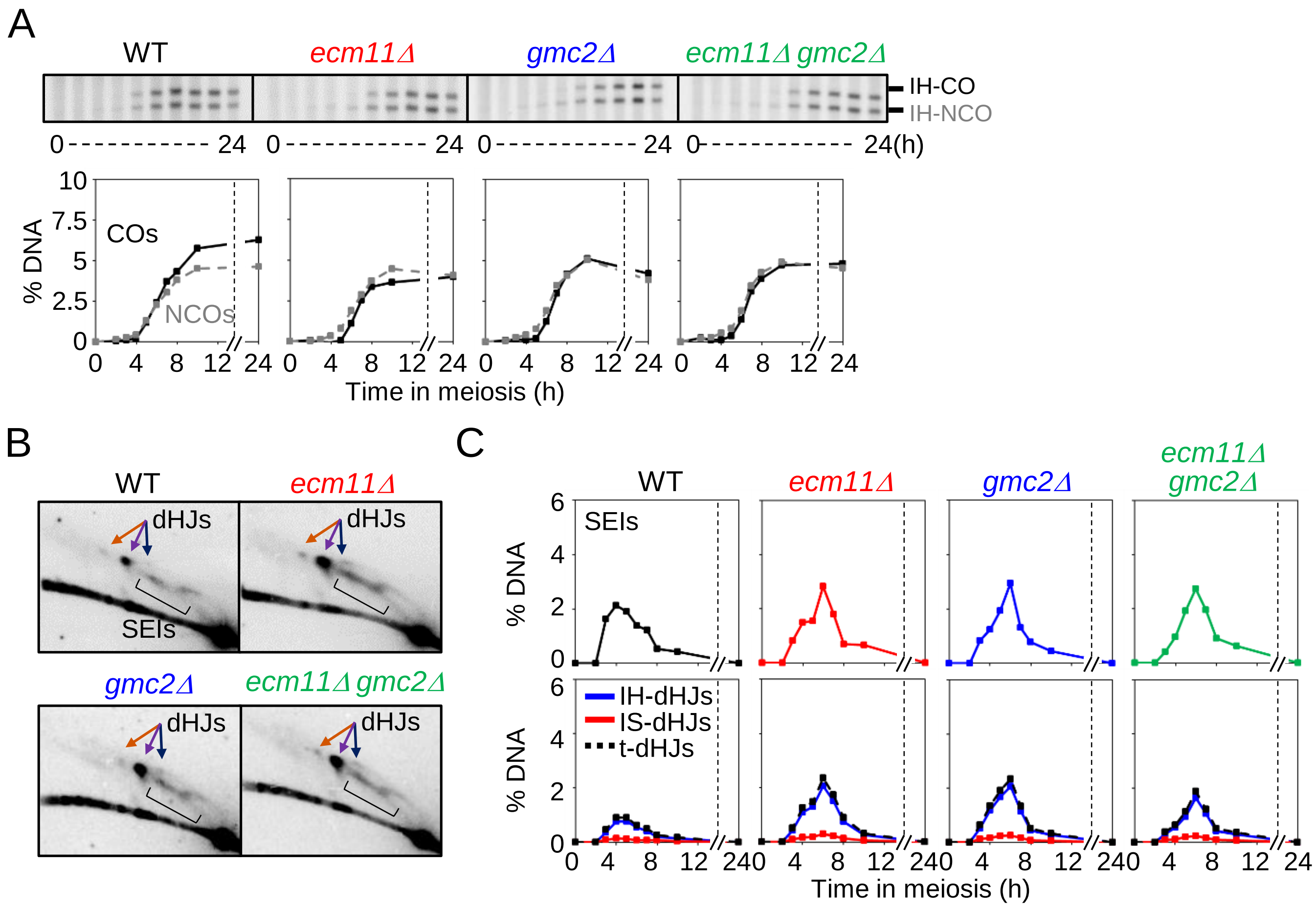

### Supplemental Figure 14

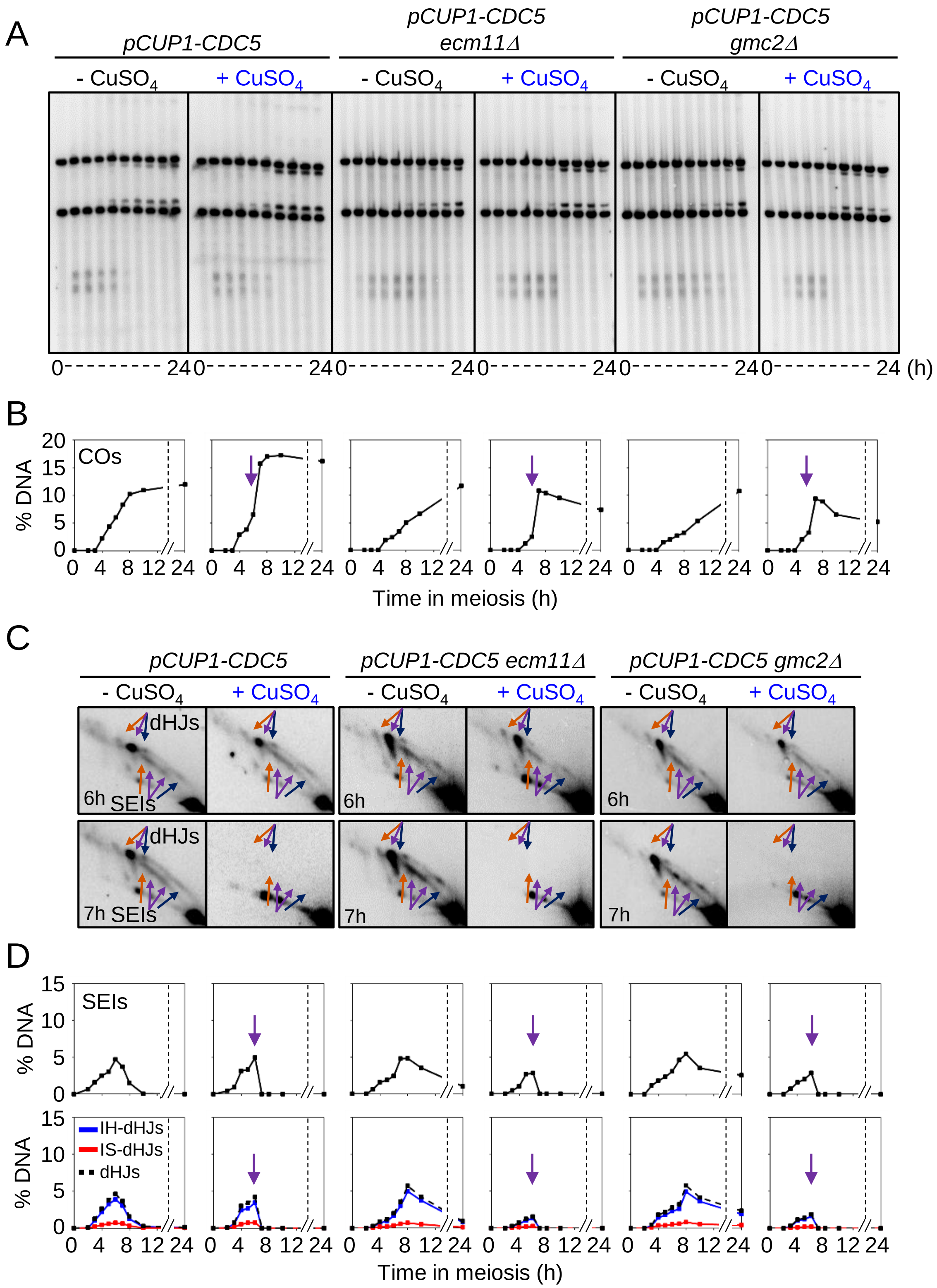
